# A 667-nucleotide sequence in the SARS-CoV-2 *nsp15* coding region promotes genome encapsidation

**DOI:** 10.64898/2026.05.29.721935

**Authors:** Shun Yao, Abhinay Gontu, Jason Atkins, Maurice Byukusenge, Padmaja Jakka, Gabriella Worwa, Jens H. Kuhn, Suresh V. Kuchipudi, Marco Archetti

## Abstract

Coronavirus genome encapsidation depends on cis-acting RNA elements that interact with viral structural proteins. While such packaging signals have been characterized in several coronaviruses, their definition in SARS-CoV-2 remains incomplete. Using synthetic defective SARS-CoV-2 genomes, we identify a 667-nucleotide region within the *nsp15* coding sequence that preferentially binds SARS-CoV-2 nucleoprotein and enhances the accumulation of defective viral genomes both in vitro and in vivo. Sequential and targeted deletion analyses further delineate candidate RNA secondary structures within this region that contribute to this enrichment. These structures show similarity to elements within the putative packaging signal of SARS-CoV but are not conserved across other coronaviruses. Together, these findings support the presence of a structured RNA element within *nsp15* that contributes to SARS-CoV-2 genome encapsidation and provide a framework for further structural and functional dissection of coronavirus packaging signals.

**IMPORTANCE:** This study identifies a 667-nt region within the SARS-CoV-2 *nsp15* coding sequence that binds nucleoprotein and promotes accumulation of defective viral genomes, revealing a previously unrecognized contributor to genome encapsidation. Mapping of candidate RNA structures within this region links SARS-CoV-2 packaging activity to conserved structural features observed in SARS-CoV, while highlighting key differences from other coronaviruses. These findings refine understanding of cis-acting packaging signals in SARS-CoV-2 and provide a foundation for further structural and functional analysis of coronavirus genome encapsidation.

GRAPHICAL ABSTRACT
A part of the *nsp15* coding sequence of SARS-CoV-2 promotes efficient transmission of defective viral genomes in vitro and in vivo. Using a sequential deletion library and targeted deletions within this region we identify RNA structures that may function as packaging signals.

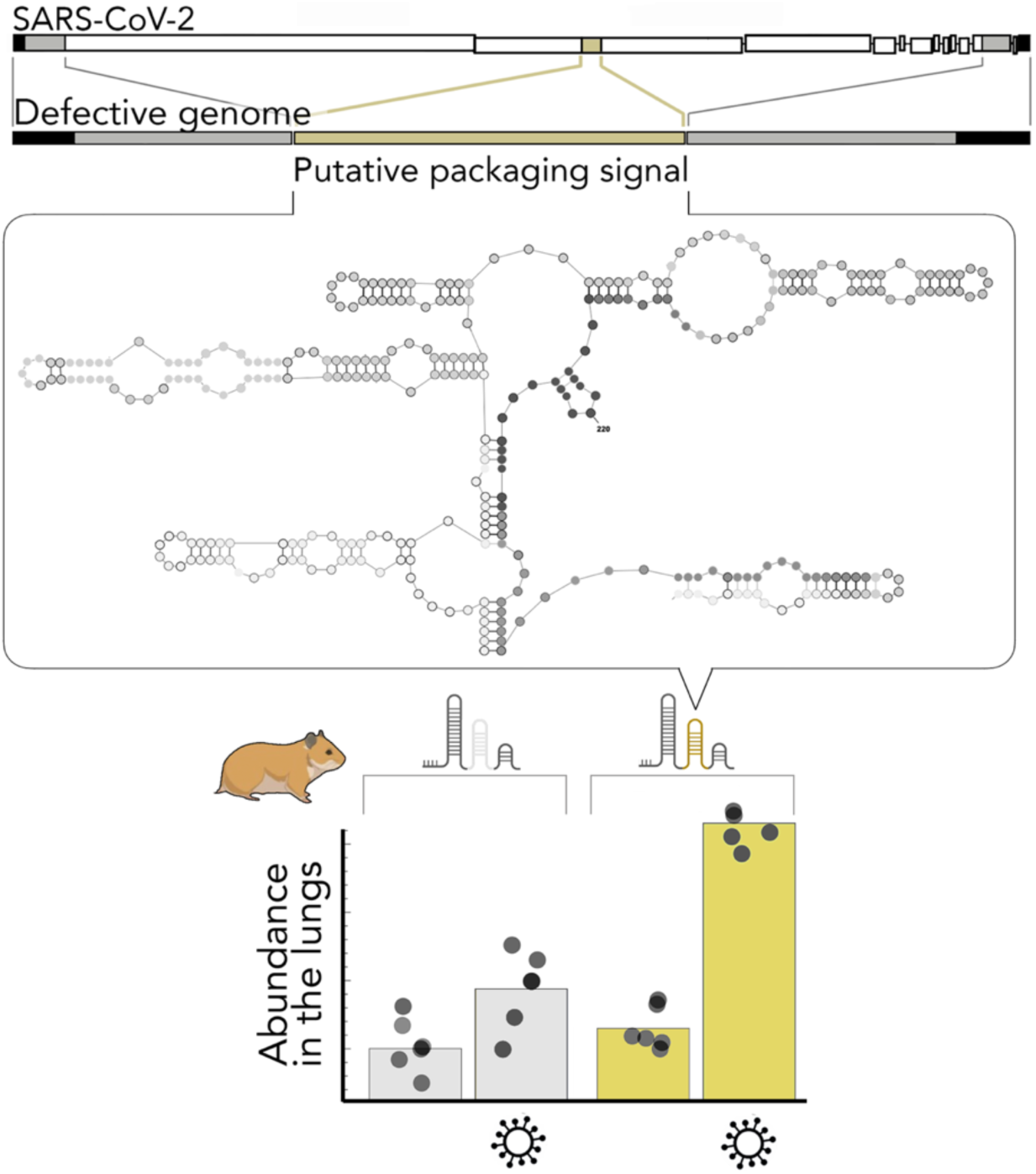

## INTRODUCTION

Viruses that spread by particle formation must ensure that their genome is incorporated into assembled virions, ideally excluding defective virus genomes and other nucleic acids present in the cell. In many positive-sense single-stranded RNA viruses (within the realm *Orthornavirae*), the efficiency and specificity of encapsidation is ensured by cis-acting RNA sequences within the virus genome. These sequences are referred to as packaging signals (“PS”) and form stable stem–loop structures that facilitate interactions with virus structural proteins.

Important examples of orthornavirans that use such packaging signals are coronavirids in the order *Nidovirales* [1–15]. Approximately half of the ∼30 kb RNA genome (*ORF1ab*) is used directly as an mRNA to produce a polyprotein that is proteolytically cleaved into nonstructural proteins (Nsps); whereas the other half is transcribed into a nested set of subgenomic RNA (sgRNA) that serve as templates for individual mRNAs that encode structural proteins. While sgRNA are sometimes packaged at low efficiency, virions contain almost exclusively genomes [1–4].

The most extensively studied coronavirus packaging signal is that of mouse hepatitis virus (MHV; species *Betacoronavirus* (*Embecovirus*) *muris*) [1–10]. The MHV packaging signal is 95 nt long and located within the *nsp15* coding sequence of *ORF1ab*, but efficient encapsidation occurs even when the packaging signal is moved within in the genome [7]. Because of the location in *ORF1ab*, the packaging signal is absent from sgRNAs and therefore sgRNAs are only rarely incorporated into coronavirus particles. The embecovirus PS is highly conserved [4,9]; indeed, the PS of bovine coronavirus (BCoV; species *Betacoronavirus* (*Embecovirus*) *gravedinis*) can be replaced with that of MHV without function loss and vice versa [10]. This PS is, however, not an absolute prerequisite for encapsidation [2], and it is not found in other betacoronaviruses, let alone in more divergent coronaviruses [4].

Different sequences within *nsp15* have been identified as potential packaging signals for Middle East respiratory syndrome coronavirus (MERS-CoV; species *Betacoronavirus* (*Merbecovirus*) *cameli*) [11] and severe acute respiratory syndrome coronavirus (SARS-CoV; species *Betacoronavirus* (*Sarbecovirus*) *pandemicum*) [12]. In transmissible gastroenteritis virus (TGEV; species *Alphacoronavirus* (*Tegacovirus*) *suis*), a packaging signal is located in the 5′-terminal region of the genome [3,4,13] and overlaps with RNA secondary structures spanning the 5′-UTR and the start of *nsp1* that contribute to genome encapsidation in alphacoronaviruses and most betacoronaviruses, but this organization differs in embecoviruses [4,14]. For infective bronchitis virus (IBV; species *Gammacoronavirus* (*Igacovirus*) *galli*), the location of a potential packaging signal was pinpointed to a 6-kb region towards the 5′ end of the genome [15].

The location and sequence of the packaging signal for severe acute respiratory syndrome coronavirus 2 (SARS-CoV-2; *Betacoronavirus* (*Sarbecovirus*) *pandemicum*) is of great interest given the impact of SARS-CoV-2 on public health [16] and the limitations of specific antiviral drugs [17] but its location remains elusive. Since packaging signals of coronaviruses reside in the replicase gene (*ORF1ab*) to avoid encapsidation of sgRNAs and must be specific enough to limit encapsidation to the SARS-CoV-2 genome, inference from cross-species homology could help determine the location of the SARS-CoV-2 PPS. However, given the diversity in packaging signal location and sequence composition across coronaviruses, the identification of the SARS-CoV-2 PPS requires direct experimental validation and systematic characterization rather than inference from cross-species homology.

The incorporation of a series of SARS-CoV-2 RNA genome fragments, each consisting of a different region of *ORF1ab*, into SARS-CoV-2-like particles indicated that a region across the *nsp15*–*nsp16* coding regions mediates encapsidation most efficiently [18]. A defective SARS-CoV-2 genome containing a different region of *nsp15* was efficiently passaged in culture in the presence of a helper full-length virus [19]. A sequence from the *nsp12*–*nsp13* coding regions enabled efficient encapsidation of a series of SARS-CoV-2 defective virus genomes [20], and this region of *nsp12*–*nsp13* is also present in naturally occurring defective genomes that accumulate during serial passages of SARS-CoV-2 [21]. In addition, a secondary structure in the 5’ end region of the genome may help encapsidation [4,14]. Finally, there is conflicting evidence as to whether defective SARS-CoV-2 genomes without these putative packaging signals can be transmitted efficiently across cells in vitro [19, 22].

We studied the encapsidation efficiency of the SARS-CoV-2 putative packaging signal (PPS) within the *nsp15* coding sequence using a combination of comparative genomic analysis, analysis of predicted secondary structures, RNA-protein binding assays, competition experiments between defective virus genomes (DVGs) with specifically designed deletions, and in vivo validation.

## RESULTS

### A 667-nt sequence encoding the middle domain of *nsp15* is a prime candidate PPS

We compared coronavirus genomes to search for candidate PPS sequences, which we expected to be variable across coronaviruses but conserved within SARS-CoV-2 variants. One such region (**Fig. 1A–B**) is within the *nsp15* coding sequence (towards the 3’ end of *ORF1ab*) and encodes the middle domain of Nsp15 (cd21167 in the NCBI’s conserved domain database [23,24]), flanked upstream by sequences encoding the conserved Nsp15 N-terminal domain and downstream by the highly conserved Nidoviral uridylate-specific endoribonuclease domain.

**Fig 1.**
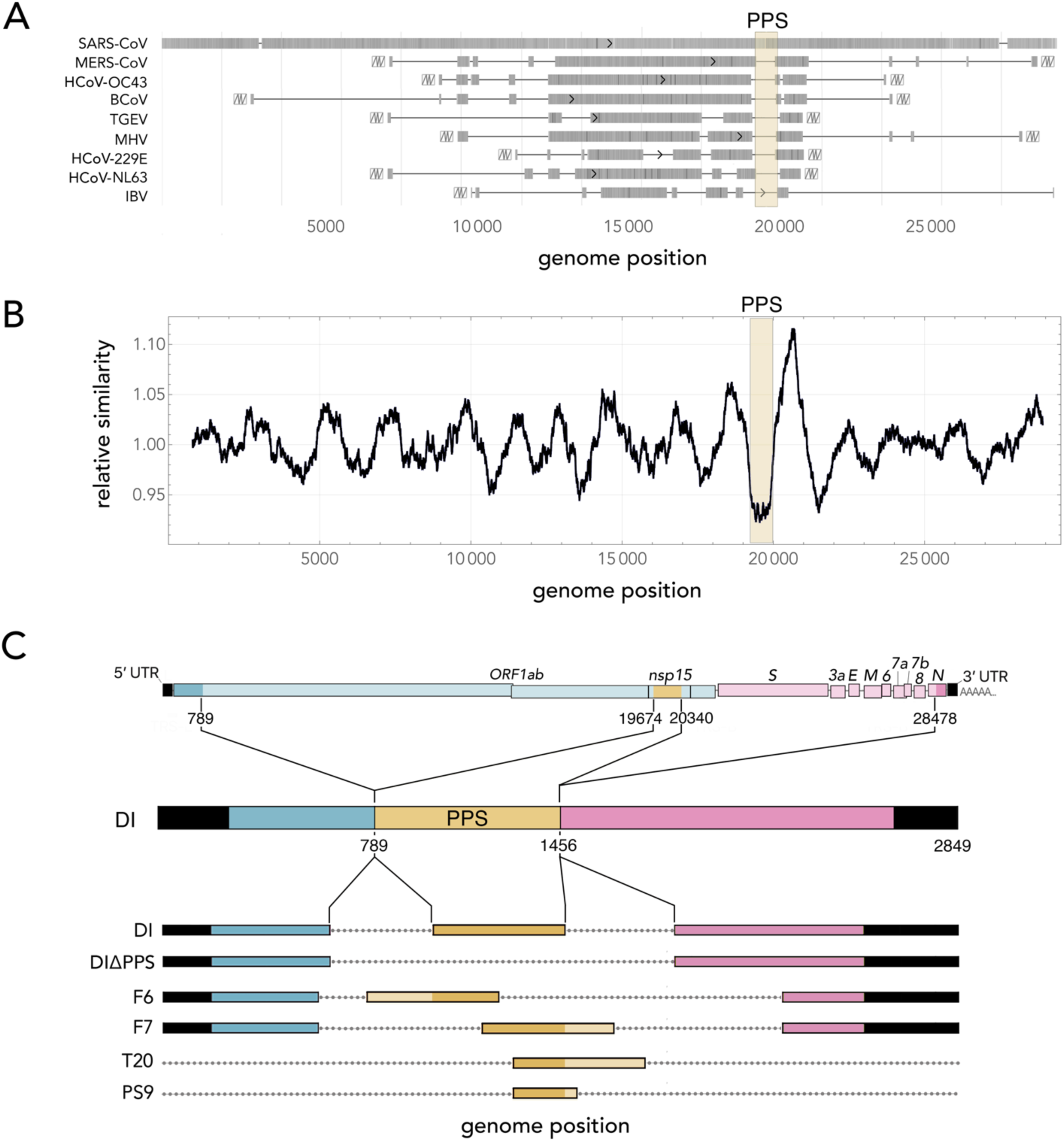
A 667-nt region of *nsp15* is a prime candidate packaging signal for SARS-CoV-2. **A**: Graphical representation of BLAST alignments among different coronavirus genomes. Darker regions denote higher alignment scores relative to the SARS-CoV-2 genome, while white boxes with zigzag lines indicate alignment boundaries. The sequence proposed as the putative packaging signal (PPS) is highlighted in yellow. **B**: Relative similarity of the SARS-CoV-2 genome with the genome of BCoV plotted as a function of genomic position, again highlighting the PPS in yellow. **C**: Genomic organization of SARS-CoV-2 and of the defective virus genomes mentioned in the text. The deletions (dotted lines) are not depicted to scale.

This part of *nsp15* corresponds to a 667-nt region (positions 19674-20340 of the SARS-CoV-2 genome; **Fig. 1C**) in the middle part of a SARS-CoV-2 synthetic defective genome that is packaged as efficiently as the full genome [19]. This region, while neither particularly conserved nor particularly variable at the protein level (**Supplementary Fig. 1**), shows substantial sequence divergence across coronaviruses (**Fig. 1A–B**; **Supplementary Fig. 2,3**). It is the most variable region of the genome relative to its flanking sequences across all evaluated coronaviruses except the closest relatives of SARS-CoV-2 (**Supplementary Fig. 2,3**). Even its homologous sequence in SARS-CoV has a limited similarity (82%).

On the other hand, the 667-nt sequence is 97.5% similar to its homologous sequence in Betacoronavirus RpYN06, which infects least horseshoe bats (*Rhinolophus pusillus* Temminck, 1834) [25,26] (the RpYN06 genome has an overall 94.5% similarity to that of SARS-CoV-2), and we did not find mutations in the 667-nt PPS across the SARS-CoV-2 variants we examined, spanning the Wuhan reference strain through Omicron lineages, a result consistent with functional constraints acting on this region.

Together, these data suggest that this 667-nt sequence within *nsp1* is a prime candidate packaging signal for SARS-CoV-2.

### The 667-nt SARS-CoV-2 PPS has high affinity for the SARS-CoV-2 nucleoprotein

We tested whether the 667-nt PPS interacts with the SARS-CoV-2 nucleoprotein (“N”) in an electrophoretic mobility shift assay (EMSA). A defective viral genome (DVG) consisting of the SARS-CoV-2 genome with the 667-nt sequence but containing two long deletions flanking it (DI in **Fig. 1C**) bound efficiently to the SARS-CoV-2 N protein and exhibited a clear mobility shift in the presence of the N protein, whereas a DVG lacking the PPS (DIΔPPS in **Fig. 1C**) displayed a mobility shift comparable to control RNA not derived from the SARS-CoV-2 genome (*EGFP* mRNA) (**Fig. 2)**. These findings indicate that the 667-nt PPS enhances RNA-N protein interaction, as would be expected for a packaging signal.

**Fig 2.**
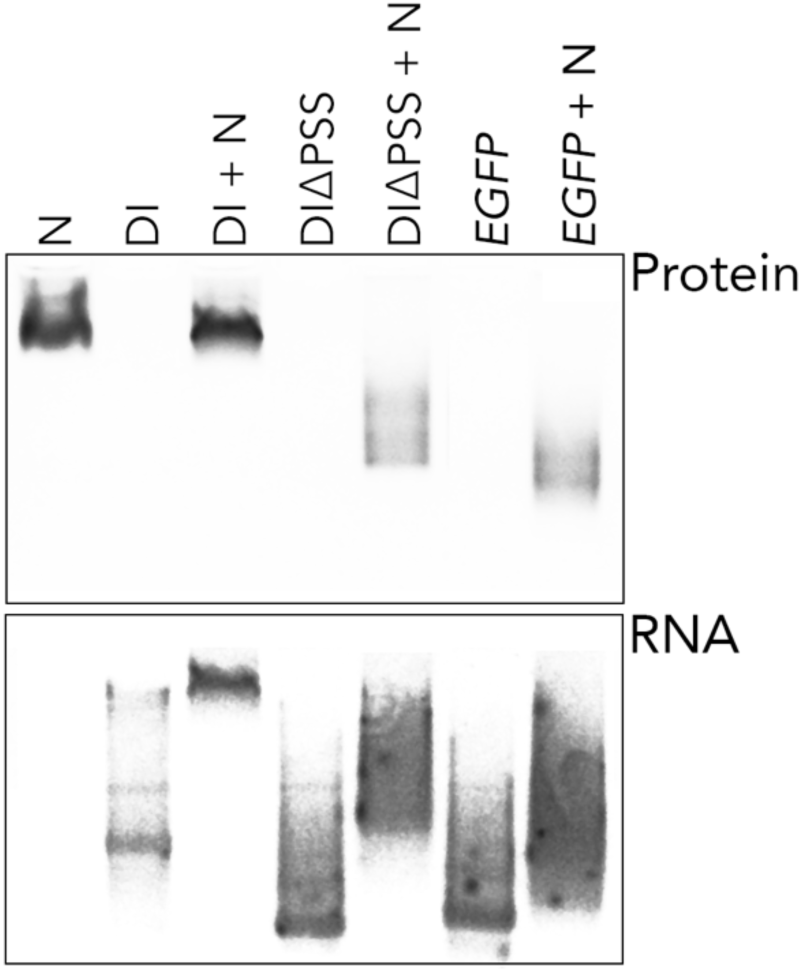
The SARS-CoV-2 putative packaging signal has high affinity for the SARS-CoV-2 nucleoprotein. Electrophoretic mobility shift assay (EMSA) of RNA SARS-CoV-2 defective genomes DI and DIΔPPS (described in **Fig 1C**) or control *EGFP* mRNA alone or complexed with SARS-CoV-2 nucleoprotein (N).

### Stepwise truncation of the 667-nt PPS defines a region required for efficient encapsidation

To identify regions of the 667-nt PPS required for efficient encapsidation, we generated systematic deletions spanning the entire PPS region within the DI DVG: a Δ3′ library with incremental 9-nt deletions from the 3′ end toward the 5′ end of the PPS, and a Δ5′ library with incremental 9-nt deletions from the 5′ end toward the 3′ end. All DI deletion variants retained identical flanking regions outside the PPS (**Fig. 3A**). The resulting libraries were pooled and transfected into A549:hACE2 cells infected with SARS-CoV-2. DI deletion variants were then quantified in the supernatant by sequencing, and relative enrichment or depletion of individual deletion variants was used to infer their packaging efficiency. This systematic analysis of DVG enrichment revealed that DI variants containing deletions spanning a region approximately 194 to 212 nucleotides downstream of the 5′ end of the PPS were significantly depleted (**Fig. 3B**). In contrast, variants with deletions outside this critical region accumulated to levels comparable to the full-length PPS control. This pattern indicates that sequences within this core region are necessary for efficient genome encapsidation.

**Fig. 3.**
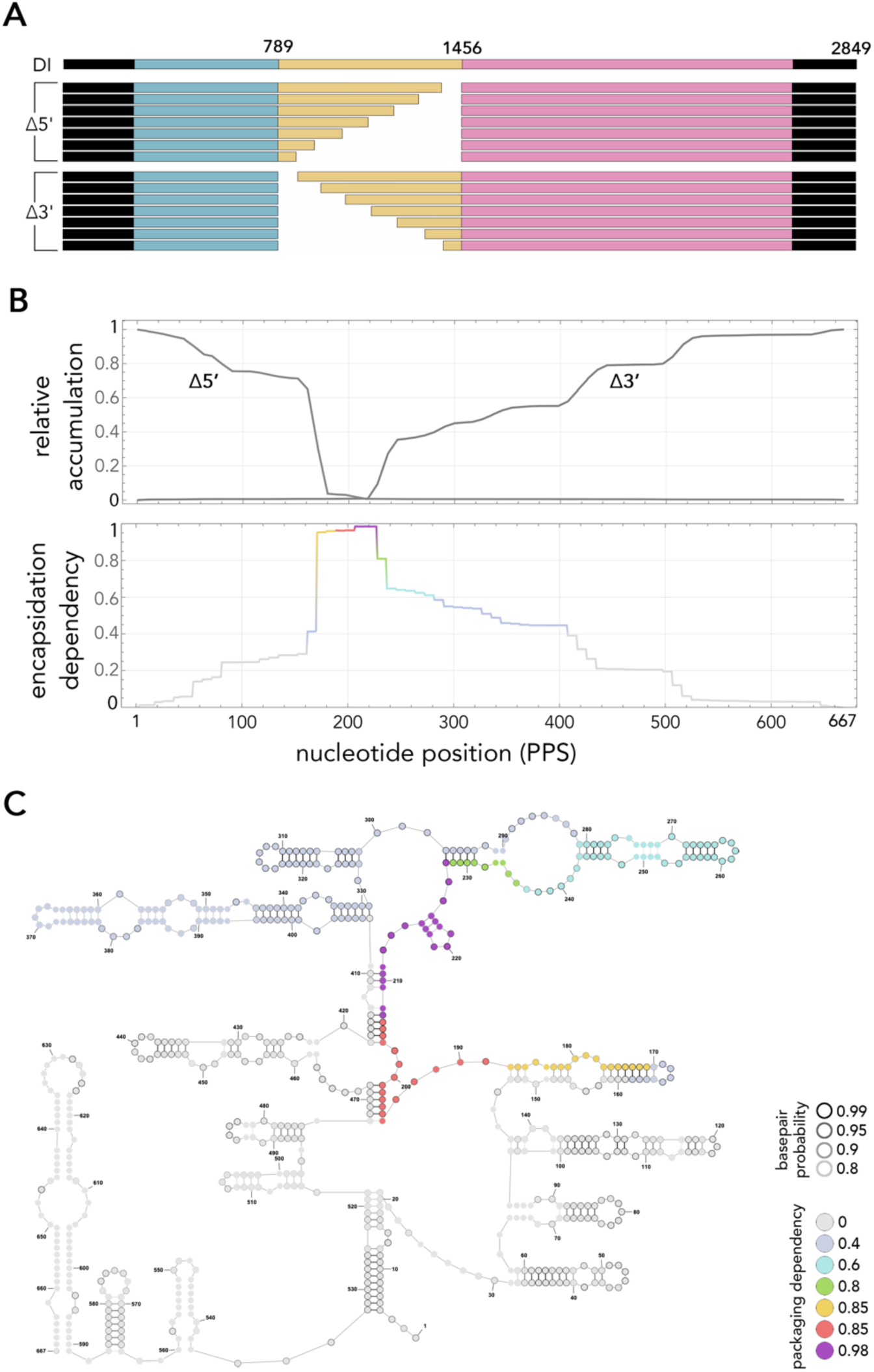
Structural analysis of the SARS-Co2 putative packaging signal reveals a core sequence and secondary structures likely important for function. **A**: Variants of the DI defective virus genomes with cumulative deletions of 9 nt starting at the 5′ (Δ5’ library) and or 3′ (Δ3’ library) ends of the 667-nt-long PPS (yellow; only seven variants per library are depicted here) were transfected into cells infected with SARS-CoV-2 and the accumulation of DI deletion variants was measured in the supernatant. **B**: Relative accumulation values and their product (encapsidation dependency) at each position of the 667-nt PPS. The packaging dependency is the product of the two values. **C**: Predicted secondary structures of the PPS and their packaging potential form panel **B**. The base pair probability is detailed in **Supplementary** Fig. 3.

### Secondary structure prediction reveals conserved stem–loops within the PPS

Since packaging signals typically depend on structured RNA elements to bind to nucleoproteins, we performed secondary structure analysis on the 667-nt PPS using thermodynamic folding algorithms (**Supplementary Fig. 4**) to identify predicted RNA secondary structures. The analysis revealed potential long-range interactions between the core 194–212 region of the PPS and a short region ∼200 nt downstream (positions 409–419 and 468–473; **Fig. 3C, Supplementary Fig.4**). Further analysis predicted that deleting nt 194–212 would disrupt most of the secondary structures of the 667-nt PPS (**Supplementary Fig. 5,6**), indicating that alterations to this region may have broad functional consequences.

**Fig. 4.**
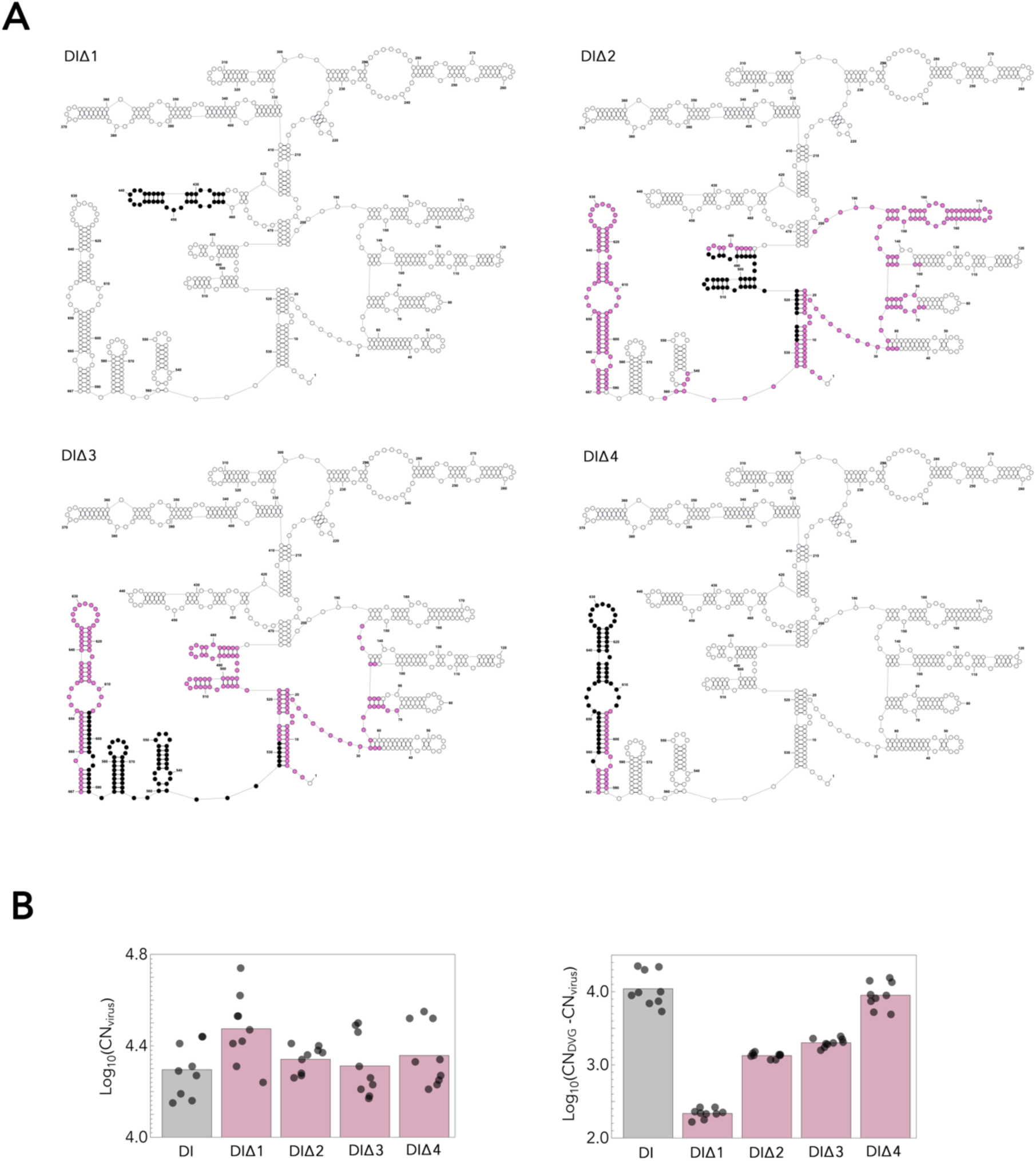
Deletion of predicted stem loops of the PPS disrupts encapsidation. **A:** Defective virus genome (DVG) DI (see Fig. 1C) was used to create four deletion variants (DIΔ1, DIΔ2, DIΔ3, and DIΔ4) with deleted nucleotides comprising specific predicted secondary structures highlighted in black. Additional secondary structures predicted to be disrupted by the deletions are highlighted in purple. **B**: Assessment of replication potential (left; none of the differences are significant: *p*>0.05; Mann-Whitney U test) and encapsidation potential (right; all the differences except that between DI and DIΔ4 are significant: *p*<0.001; Mann-Whitney U test) of the DVG variants.

Next, we compared the predicted secondary structure of the PPS with that of its homolog in SARS-CoV (PS580 [12]; nt 19605–20270 of the SARS-CoV genome). Interestingly, five long stem loops and two shorter ones are shared between the two virus genomes (**Supplementary Fig. 7,8**) including the two long stem loops predicted to be formed by nt 227–327 of the 667-nt PPS of SARS-CoV-2, i.e., the region with the highest encapsidation potential (**Fig. 3C**). One of the predicted stem loops in the 667-nt PPS (SL5 in **Supplementary Fig. 7, 8**) is the central part of the core of the SARS-CoV putative packaging signal PS580 (PS63, nt 19888–19950 [12]). One of these (SL5 in **Supplementary** and SL5, is found only in sarbecoviruses, among the coronaviruses we analysed here (**Supplementary Fig. 3**).

A stronger structural similarity is predicted with the homologous sequence of bat coronavirus RpYN06: most of the secondary structures predicted for the 667-nt PPS of SARS-CoV-2, including the long-range interactions, are also predicted to occur in this virus (**Supplementary Fig. 9**). The only secondary structures not predicted for bat coronavirus RpYN06 are the ones towards the 3’ end (nucleotides 549-667) of the 667-nt PPS. RaTG13, another sarbecovirus, has no structural equivalent of SL2, despite high nucleotide similarity (**Supplementary Fig. 3**) but it has a more extensive SL4 structure (**Supplementary Fig. 10)**. None of the secondary structures of the 667-nt PPS are predicted to occur in the homologous regions of other coronaviruses analysed here, including MERS-CoV (**Supplementary Fig. 11**) and MHV (**Supplementary Fig. 12)**, suggesting that the secondary structures predicted for the 667-nt PPS evolved within the sarbecovirus clade.

Finally, we analyzed the secondary structures of DVGs used in previous studies of SARS-CoV-2 packaging signals [18,20] to understand why this 667-nt PPS was not described in two previous studies. Our analysis shows that the DVGs used in these two studies disrupt the 667-nt PPS at position 203 or 274 [20] or at positions 210 and 406 [18], preserving none or few of the secondary structures identified here (**Supplementary Fig. 13-17**).

### Deletion of predicted stem loops of the PPS reduces encapsidation efficiency

To test the importance of the secondary structures identified above and refine the location of the packaging signal, we designed and synthetized four DVG variants (DIΔ1, DIΔ2, DIΔ3, DIΔ4) based on DI DVGs with deletions that disrupt specific predicted secondary structures (**Fig. 4A and Supplementary Fig. 18-21**). We then transfected these DI variants separately into A549:hACE2 cells infected with SARS-CoV-2, and measured DVG RNA levels both in infected cells and in the viral supernatant. The deletion mutants were equally efficient replicators as DI based on the amounts of DVGs variants and viral RNA measured in cell extracts (**Fig. 4B**; none of the differences are significant: *p*>0.05; Mann-Whitney U test). We then compared the ratio of DVG RNA to SARS-CoV-2 genomic RNA in the supernatants, a measure of the encapsidation potential of the putative packaging signal with and without the deletions. DVG variants DIΔ1, DIΔ2, DIΔ3 exhibited significantly reduced DVG-to-genomic RNA ratios in the supernatant compared to DI, whereas DIΔ4 showed no significant reduction in packaging efficiency (**Fig. 4C**; all the differences except that between DI and DIΔ4 are significant: *p*<0.001; Mann-Whitney U test).

This suggests that a predicted secondary structure at the 3′ end of the 667-nt PPS (nt 589–667), which is fully disrupted by the deletion in DVG variant DIΔ4, does not affect encapsidation, which is consistent with its low overall base-pair probability. Given that DVG variant DIΔ3’s encapsidation efficiency was somewhat reduced, its two stem loops at 537–587 and at 475–516, and the long-distance pairing of nt 518–533 with nt 3–21 may be involved encapsidation. Other secondary structures at nt 31–97 were only minimally affected by the deletion. DVG DIΔ2’s encapsidation efficiency was further reduced; hence, the two predicted stem loops at 537–587 identified above, which are not affected by this deletion, are likely less important for encapsidation than the ones at 475–516, which remain affected by this deletion. Deletion of an additional stem loop at nt 147–188 additional structural changes compared to DIΔ3 affect other stem loops in nt 31–143 only marginally. Disruption of the predicted stem loop at 425–459 in DIΔ1, which was not changed otherwise, led to the most dramatic reduction in encapsidation efficiency. Given that this is the only predicted secondary structure disrupted in DIΔ1, and that this secondary structure was not disrupted in the other variants, this stem loop likely is a key determinant of encapsidation.

### The PPS enhances defective virus genome accumulation *in vivo*

To assess the importance of the 667-nt PPS in infections in vivo, we administered nanoparticles containing DVG variant DI RNA, DIΔPPS RNA, or control *EGFP* mRNA to golden hamsters previously exposed to SARS-CoV-2 or mock control and then measured the amount of the three RNAs in the lungs 4 days after administration (**Fig. 5**). In uninfected animals all RNAs reached the lungs in small amounts (less than the administration dose). In infected animals, *EGFP* RNA did not increase compared the amount in non-infected animals; DIΔPPS was one order of magnitude more abundant in infected hamsters than in uninfected hamsters, and one order of magnitude more abundant than EGFP mRNA, but almost three orders of magnitude less abundant than DI RNA (**Fig. 5**; Mann-Whitney U test, *p*<0.0001). These findings indicate that the PPS element dramatically enhanced DI RNA accumulation in infected lungs.

**Fig. 5.**
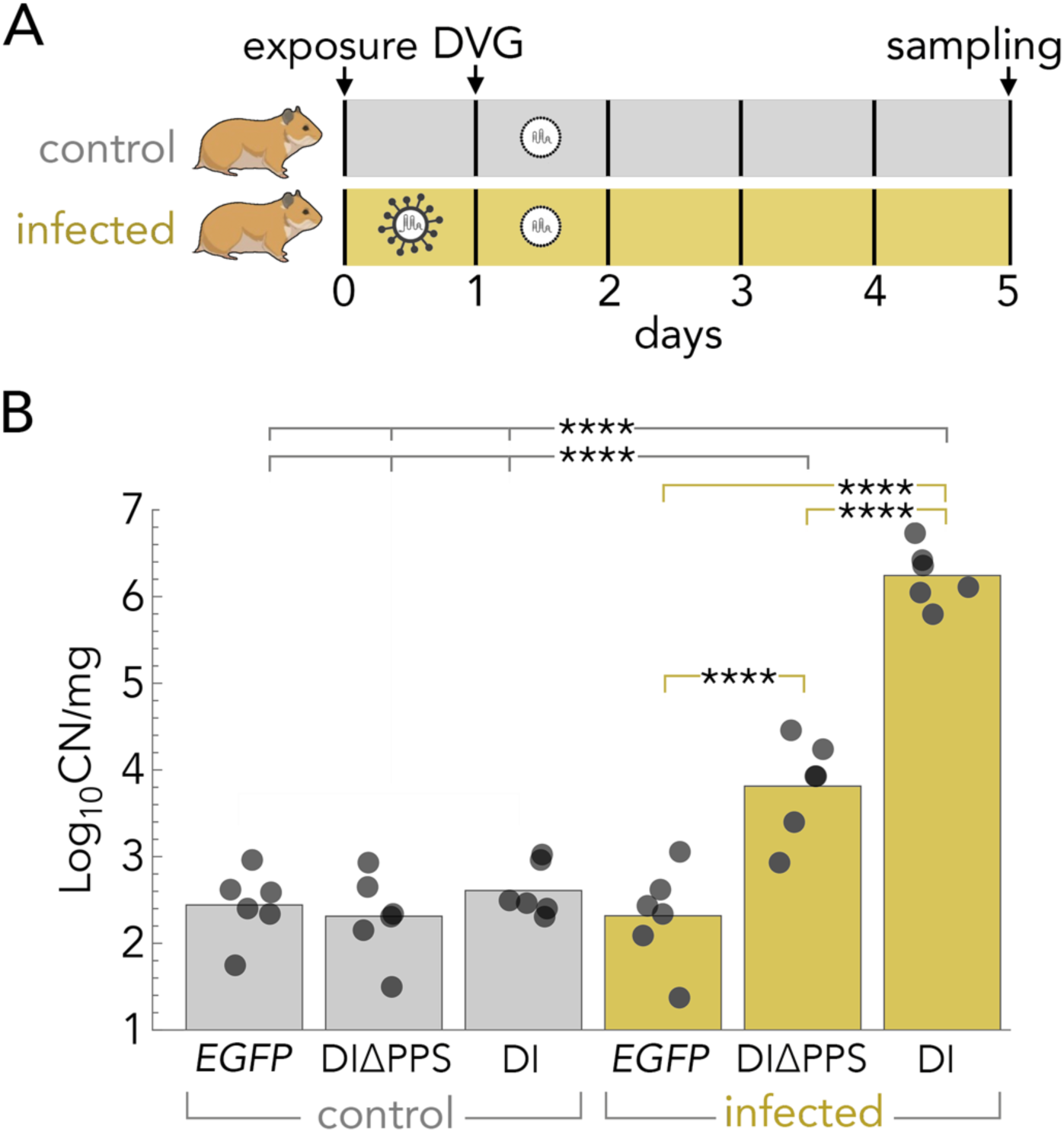
The 667-nt PPS increases the amount of defective virus genomes in vivo. **A:** Golden hamsters exposed to SARS-CoV-2 or mock control on Day 0 were inoculated intranasally on day 1 with nanoparticles carrying SARS-CoV-2 defective virus genome (DVG) DI (containing the complete 667-nt putative packaging signal), DIΔPPS (DI without the 667-nt sequence), or control *EGFP* mRNA. Lung samples were collected on Day 5 for RNA quantification. **B**: The amount of RNA, expressed as Log_10_ of the copy number (CN) per mg of lung samples. **** *p*<0.0001, Mann-Whitney U test.

## DISCUSSION

Taken together, our data indicate that a 667 nt long sequence in the middle domain of the *nsp15* coding sequence (positions 19,674–20,340) promotes efficient genome packaging through RNA structural features and enhanced interaction with nucleocapsid protein, while exhibiting evolutionary conservation in genomic position but substantial nucleotide sequence and structural divergence among related coronaviruses.

Efficient genome packaging in coronaviruses depends on interactions between viral RNA and the N protein. Consistent with this requirement, RNA constructs containing the 667-nt PPS exhibit enhanced mobility shifts in EMSA, indicating greater association with N protein compared to control sequences lacking the element. This observation aligns with previous studies demonstrating that RNA-N protein interactions are central to encapsidation in SARS-CoV and other coronaviruses [3,8,12,27,28]. However, our EMSA approach does not provide quantitative binding parameters, cannot distinguish between sequence-specific recognition and recognition of secondary structures, and does not reveal the molecular basis of this interaction. Addressing these questions will require, respectively, quantitative binding assays, targeted mutational and compensatory structure-disruption analyses, and structural biology approaches to identify molecular contacts critical for binding. Together, these findings nonetheless support a functional role for the 667-nt element in promoting N protein association and highlight it as a key cis-acting determinant of coronavirus genome packaging.

Building on this evidence of enhanced RNA–N protein interaction, we examined the evolutionary context and structural conservation of the 667-nt PPS across coronaviruses. The 667-nt PPS has a high nucleotide divergence across species, while residing in an otherwise conserved region of *ORF1ab*. It overlaps substantially with the SARS-CoV putative packaging signal PS580 (nt 19715–20294) [12], and despite only 82% predicted to occur also in PS580. Indeed, one of the stem loops in the 667-nt PPS corresponds to PS580 core (PS63 [12]; nt 19888–19950). Putative packaging signals have been identified in homologous *nsp15* regions in both sarbecoviruses [12] and merbecoviruses (MERS-CoV [11]), suggesting a shared genomic location even though their sequences and structures are free to evolve and consistent with the lack of the embecovirus PS in the sarbecovirus *nsp15* [3,4]. The genomic location of the 667-nt PPS within *nsp15*, flanked by recombination breakpoints [26], further suggests the hypothesis that packaging signals could be acquired through recombination events. Together, these findings suggest that the 667-nt PPS represents a structurally conserved yet evolutionarily flexible packaging module, highlighting its potential importance in shaping coronavirus genome evolution.

To define the functional subregions within the 667-nt PPS more precisely, we next performed systematic deletion mapping using complementary 5′ and 3′ truncation libraries identified approximately 220 nucleotides downstream of the 5′ end of the PPS, where deletion variants were progressively depleted in co-infection with wild-type virus. An important consideration for interpreting this is that our deletion library used a competitive format forcing DVGs lacking critical packaging signal elements to compete directly with DVGs containing a fully functional part of the 667-nt PPS. In non-competitive scenarios in which all DVGs lack that sequence, DVGs may persist despite reduced encapsidation efficiency due to diminished competition for the packaging machinery. In other words, the 667-nt PPS and its core region may enhance, rather than being absolutely required for, encapsidation.

To further characterize this region, we generated targeted deletion mutants designed to disrupt specific predicted secondary structures. These experiments revealed a hierarchical organization of functional elements: disruption of the stem-loop at positions 425-459 (DIΔ1) produced the most substantial reduction in packaging efficiency (∼70% decrease compared to intact PPS), indicating that this element plays a major functional role. Sequential analysis of additional mutants showed progressively greater defects: DIΔ2 and DIΔ3, which disrupt stem-loops at positions 147-188 and 475-516 respectively, resulted in smaller but measurable cumulative reductions in packaging. In contrast, disruption of the 3′-terminal stem-loops (positions 589-667) in DIΔ4 did not significantly impair packaging efficiency, indicating that these distal structures are functionally dispensable or redundant.

A caveat with these results is that the stem-loops at positions 147–188, 425–459, and 475–516, which we identified as functionally important, are located proximal to, but not precisely within, the region that had the highest packaging dependency in our stepwise deletion assay (positions 194–212). This apparent discrepancy may arise from several factors. First, the 9-nucleotide deletion increments used in our library approach may lack sufficient resolution to pinpoint the most critical elements — individual nucleotide substitution mutagenesis would be required to pinpoint precisely which nucleotides are critical. Second, our stepwise deletion approach measured the downstream effects of deletions on encapsidation rather than directly assessing the intrinsic encapsidation potential of specific sequences — effects that may operate indirectly through adjacent secondary structures. Finally, the possibility remains that additional genomic sequences beyond the identified 667-nt PPS sequence contribute to overall encapsidation.

Overall, these results support a model in which multiple RNA secondary structures cooperatively contribute to packaging efficiency, with the stem-loop at 425-459 playing a predominant role. Secondary structure predictions further indicate that this central region contains multiple stem-loop elements and long-range RNA interactions that may facilitate nucleocapsid protein binding. Our conclusions are based on defective viral genomes rather than full-length SARS-CoV-2 genomes, which may not fully recapitulate native packaging dynamics. Although DVG systems are widely used to study coronavirus packaging, differences in genome size, RNA secondary structure, replication kinetics, and competition with wild-type virus could influence packaging and assembly behavior relative to intact viral genomes.

In vivo validation in hamsters demonstrated that DI RNA containing the intact 667-nt packaging element accumulated more efficiently in infected lungs than DI RNA lacking the element. Although the enhanced accumulation could reflect increased packaging, replication, stability, or spread, the consistent and dramatic effect observed across both in vitro and in vivo systems provides strong evidence that this element plays a major functional role in the SARS-CoV-2 infectious cycle.

Previous studies had pointed to different putative packaging signals. Minigenomes comprising the utmost 5’ 742 nt and terminal 3’ 605 nt, flanking a region derived from the *nsp12*–*nsp13* coding sequence, are efficiently packaged, and altering the secondary structure of this region affected packaging efficiency [20]. Conclusions were based on systematic analysis of DVGs designated F1–F7 (shown for reference in **Fig. 1C**), ranging from 1–1.7 kb in length and spanning the *nsp7*–*nsp16* coding regions. However, this fragmentation approach inadvertently bisected the 667-nt PPS we studied here: DVG F6 terminates at position 274 of our PPS, whereas DVG F7 begins at position 203, thereby disrupting the secondary structures we identified as essential for encapsidation, potentially explaining why Terasaki et al. [20] did not detect the 667-nt PPS we describe here. The *nsp12*–*nsp13* region encapsidated, suggesting that multiple packaging signals may exist within the SARS-CoV-2 genome.

Similarly, Syed et al. [18] examined encapsidation of SARS-CoV-2 RNA fragments spanning different *ORF1ab* regions in a virion-like particle assay and identified a region in *nsp15*–*nsp16* (DVG T20, positions 20,080–22,222; see **Fig. 1C**) that was packed with greater efficiency than the *nsp12* region identified by Terasaki et al. [20]. However, a direct comparison with our results is challenging because that experimental design also fragmented the 667-nt PPS at critical positions. T20 and its associated PS9 region (see **Fig. 1C**) begin at position 406 of the 667-nt PPS, whereas another tested region (PS576) terminates at position 210, both disrupting the secondary structures we identified as functionally important.

The relative encapsidation efficiencies of the putative packaging signals from all these studies are difficult to compare. Notably, F7 [20] encompasses PS9 [18]; additionally, Syed et al. [18] observed that some fragments were packaged less efficiently than shorter fragments they include, suggesting that encapsidation determinants involve complex structural relationships beyond simple two-dimensional secondary structure predictions. These observations indicate that systematic fine-scale analysis is required to identify the critical secondary structures governing encapsidation efficiency. Given the evidence for packaging activity from *nsp12*–*nsp13* [20], a different region of *nsp15*–*nsp16* [18], and potentially the 5’ UTR [3,4], SARS-CoV-2 and other coronaviruses may employ multiple genomic elements to improve encapsidation efficiency.

Our work identified a strong packaging signal for SARS-CoV-2 and provides a framework for understanding coronavirus encapsidation specificity more broadly. The identification of critical secondary structures within the packaging signal offers potential targets for antiviral development aimed at disrupting coronavirus genome encapsidation. Future structural studies examining the molecular contacts between packaging signal elements and the nucleoprotein, a finer dissection of the position of the most important secondary structures, and a direct comparison of all hypothesized packaging signals, will further illuminate the mechanistic basis of selective coronavirus genome encapsidation and clarify which sequences drive efficient encapsidation.

### Viruses and genome sequences

The viruses used for analysis and their RefSeq/GenBank accession numbers are:

- SARS-CoV-2 (Severe acute respiratory syndrome coronavirus 2, Wuhan-Hu-1): NC_045512.2
- SARS-CoV (Severe acute respiratory syndrome coronavirus, Tor2): NC_004718.3
- MERS-CoV (Middle East respiratory syndrome-related coronavirus, HCoV-EMC/2012): NC_019843.3
- HCoV-NL63 (Human coronavirus NL63, Amsterdam I): NC_005831.2
- HCoV-229E (Human coronavirus 229E, inf-1): NC_002645.1
- HCoV-OC43 (Human coronavirus OC43, ATCC VR-759): NC_006213.1
- BCoV (Bovine coronavirus, BCoV-ENT): NC_003045.1
- TGEV (Transmissible gastroenteritis virus, PUR46-MAD): NC_038861.1
- MHV (Murine hepatitis virus, A59): NC_048217.1
- IBV (Infectious bronchitis virus, Ind-TN92-03): NC_048213.1
- RpYN06 (Betacoronavirus sp. RpYN06, bat/Yunnan/RpYN06/2020): MZ081381.1
- RaTG13 (Bat coronavirus RaTG13): MN996532.2

The SARS-CoV-2 variants used to detect mutations in the 667-nt PPS are

- Alpha (B.1.1.7), GenBank: MW422255.1
- Beta (B.1.351), GenBank: MW981442.1
- Gamma (P.1), GenBank: MZ169911.1
- Delta (B.1.617.2), GenBank: OX003053.1
- Omicron (BA.2), GenBank: OM095411.1

### Defective virus genome designations

The designations for SARS-CoV-2 defective virus genomes (DVGs) and the SARS-CoV-2 genome nucleotides they contain are:

- DI [19]: 1–789, 19674–20340, 28478–29870
- DIΔPPS: 1–789, 28478–29870
- DIΔ1: 1–789, 19674–20098, 20134–20340, 28478–29870
- DIΔ2: 1–789, 19674–20158, 20202–20340, 28478–29870
- DIΔ3: 1–789, 19674–20201, 20281–20340, 28478–29870
- DIΔ4: 1–789, 19674–20281, 20335–20340, 28478–29870
- F6 [20]: 1–742, 18406–19947, 29265–29870
- F7 [20]: 1–742, 19877–21556, 29265–29870
- T20 [18]: 20080–22222
- PS9 [18]: 20080–21171

A visual presentation of these DVGs is shown in **Fig. 1C**.

Genome-level sequence similarities reported as graphical alignment in Fig. 1 were obtained using the National Center for Biotechnology Information (NCBI) BLAST suite [29] at https://blast.ncbi.nlm.nih.gov/Blast.cgi using blastn default nucleotide scoring parameters. The absolute similarity for each position in Supplementary Fig. 1 and 2 is the percentage of nucleotides that do not differ between aligned homologous sequences, calculated across the 50 amino acid (for protein similarity) or 800 nt (for nucleotide similarity) around that position. Local similarity is the absolute similarity of that position divided by absolute similarity 800 nt or 50 amino acids upstream and downstream, that is, the similarity of those 800 nt or 50 amino acid sequence compared to its flanking sequences of the same length. Sequence alignment for *nsp15* was performed using MAFFT version 7 [30] using default parameters.

### Stepwise deletion libraries

Custom DI RNA deletion libraries were designed in-house and manufactured by Thermo Fisher Scientific (GeneArt Gene Synthesis and Variant Libraries service). Two complementary libraries were generated to systematically truncate the PPS at either terminus. For the Δ3’ deletion library, we specified a series of DI RNA variants containing stepwise cumulative 9-nt deletions extending from the 3’ end of the PPS toward its 5’ end. The resulting library comprised a set of constructs in which 9, 18, 27, and subsequent 9-nt increments were removed in a stepwise manner along the 3’→5’ direction. For the Δ5’ deletion library, an analogous design strategy was applied to generate DI RNA variants with cumulative 9-nt deletions originating at the 5’ end of the PPS and extending toward the 3’ end. This library therefore contained constructs with 9, 18, 27, and further 9-nt deletions in the 5’→3’ direction. The two regions flanking the PPS were the same for all DVG variants. Thermo Fisher synthesized all variant sequences according to these design specifications and delivered them as sequence-verified DNA constructs that we used to synthetize RNA by in vitro transcription as described above.

### RNA and DVGs synthesis

DVGs RNAs and control *EGFP* mRNA were synthesized by *in vitro* transcription using the HiScribe High Yield RNA Synthesis Kit (New England Biolabs #E2040) and custom-made plasmids manufactured by Thermo Fisher Scientific (GeneArt Gene Synthesis and Variant Libraries service) for all DVGs and pGEM4Z-EGFP (Addgene #183475, a gift from Chantal Pichon) for *EGFP* mRNA. Following transcription, RNAs were purified using a spin RNA cleanup kit (New England Biolabs #T2040), RNA integrity and concentrations were assessed by automated electrophoretic separation and detection using an Agilent TapeStation system. Purified RNAs were stored at −80°C.

### Electrophoretic mobility shift assay (EMSA)

His-tagged nucleoprotein (N; Accession #YP_009724397.2; R&D Systems #10999-CV-100) was resuspended in phosphate-buffered saline (PBS) at 500 μg/ml. To perform electrophoretic mobility shift assays (EMSAs), 2 µg of DVG RNA and 5 µg of N protein were suspended alone or in combination in total volumes of 12 µl and then mixed after addition of 2 µl binding buffer (750 mM KCl, 0.5 mM dithiothreitol, 0.5 mM EDTA, 50 mM Tris, pH 7.4; Thermo Fisher Scientific #E33075, Component E) and left to incubate at room temperature for 20 min. 2.5 µl of EMSA gel loading solution (Thermo Fisher Scientific #E33075, Component D) was added before loading the samples on a 1% agarose gel. The gel was then subjected to electrophoresis. To prevent interference of the staining dyes with RNA-protein interactions and migration during electrophoresis, we stained the RNA and proteins after the entire gel-shift was completed [31]. RNA was stained in SYBR green solution made from 5 μL of 10,000x SYBRgreen (Thermo Fisher #E33075, Component D) in 50 mL of TAE buffer (40 mM Tris-acetate, 1 mM EDTA, pH ∼8.0), for 20 minutes in the dark with continuous agitation on an orbital shaker at 50 rpm, followed by washing twice in 150 mL of dH_2_O for ∼10 s each. RNA was visualized in-gel on a Chemiluminescent Imaging System (Azure Biosystems #Azure300) using excitation/emission settings of 255, 495/520 nm. N protein was stained by then covering the gel with SYPRO Ruby EMSA protein gel stain (Thermo Fisher Scientific#E33075, Component D) mixed with TCA (Thermo Fisher Scientific#E33075, Component C)] for 8 h in the dark with continuous agitation on an orbital shaker at 50 rpm followed by washing twice in 150 mL of dH_2_O for ∼10 s each, washing in 10% methanol, 7% acetic acid for 60 min, and washing twice in 150 ml of dH_2_O for ∼10 s each. N protein was visualized in-gel on the same system used for RNA visualization using the UV302 channel using excitation/emission settings of 280, 450/610 nm.

### Secondary structure prediction and analyses

Secondary structure prediction and analyses were performed using the RNAstructure [32] webserver (https://rna.urmc.rochester.edu/RNAstructureWeb/). Base-pair probabilities were calculated using the Partition software; sequences were folded based on their lowest free energy conformation using the Fold software; and the resulting structures were compared using the CircleCompare software. Default parameters were used for all analyses.

### In vitro infections

Vero cells derived from *Chlorocebus sabaeus* kidney epithelium (BEI Resources #NR-59618) and human lung adenocarcinoma A549 cells expressing human angiotensin converting enzyme 2 (hACE2), the receptor of SARS-CoV-2 (A549:hACE2 cells; BEI Resources #NR-53821), were maintained under standard *in vitro* conditions in Dulbecco’s Modified Eagle Medium (DMEM) supplemented with 10% fetal bovine serum (FBS). Cell lines were incubated at 37°C in a humidified atmosphere containing 5% CO₂ and were routinely monitored for morphology and confluency. Cells were passaged using enzymatic dissociation when reaching 70–90% confluency and were periodically tested to ensure the absence of mycoplasma contamination. For experimental procedures, cells were seeded at appropriate densities to achieve optimal growth and transfection efficiency.

SARS-CoV-2 (Isolate USA-WA1/2020, BEI Resources #NR-52281 (which differs only by a small number of nucleotide substitutions from isolate Wuhan-Hu-1) virus stocks were prepared by propagating in Vero cells. All manipulations involving live SARS-CoV-2 were conducted in biosafety level 3 (BSL-3) facilities of the Eva J. Pell Laboratory for Advanced Biological Research at the Pennsylvania State University, in compliance with the Institutional Biosafety Committee (IBC), ensuring safety protocols in accordance with federal requirements.

At the indicated timepoints, infected cells were washed twice with PBS. 1 mL of TRIzol (Thermo Fisher Scientific, #15596026) were added and incubated at room temperature for 5 min, followed by freezing at −80°C until further analysis. RNA was isolated from cell extracts or cell supernatants according to the protocol of the MagMAX Viral/Pathogen Nucleic Acid Isolation Kit (Thermo Fisher Scientific #A42352). Extracted RNA quality was assessed using a NanoDrop spectrophotometer (Thermo Fisher Scientific) to determine the absorbance ratios at 260/280 nm and 260/230 nm, and RNA integrity was further evaluated using an Agilent 2100 Bioanalyzer (Agilent Technologies). Only samples with an RNA integrity number (RIN) >7.0 were used for downstream applications.

### RNA delivery in vitro and in vivo

For cell transfection, 500 ng of each RNA were delivered by a cationic lipid–based transfection reagent (MessengerMax; Thermo Fisher Scientific #LMRNA015) prepared according to the manufacturer’s recommendations. 10 μl of RNA solution and 5 μl of MessengerMAX were mixed to support the formation of lipid–RNA complexes for 10 min, then diluted in 125 μl of Opti-MEM (Gibco #31985070), and then delivered to Vero cells that had been exposed 24 h earlier to helper SARS-CoV-2 at a multiplicity of infection (MOI) of 0.1 in 6-well plates at 70–90% confluency.

For in vivo delivery, dimethyl sulfoxide (DMSO) was added to 1 mg of D90−118 hyperbranched poly(beta-amino ester) (hPBAE) [33] and sonicated until dissolved (10-60 s.); the same volume of sodium acetate (0.1 M, pH 5.2) was then added and the solution sonicated again a few seconds to one min, yielding an hPBAE solution at 10 μg/μL. RNA was added to hPBAE at a 40:1 (w/w) hPBAE:RNA weight ratio, and mixed by pipetting 10 times.

### Quantification of RNA

Quantification was performed by one-step RT-qPCR using dual-labeled hydrolysis (TaqMan) probes specific for each DVG in combination with DVG-specific primers on a StepOne Plus Real-Time PCR System (Applied Biosystems). All reactions were run in triplicates, with no-template controls included. RNA in supernatants was quantified using a standard curve generated from serial dilutions of RNA from the products of a 20 µL reaction containing 1 µg RNA, 10 µL of FastVirus master mix (Applied Biosystems # 4444434), 0.2 µM of TaqMan probe, and 0.5 µM of primers at the following conditions: 50°C for 10 min, 95 °C for 2 min, followed by 40 cycles of 95 °C for 15 s and 60 °C for 1 min. For RNA in cell extracts, results were normalized using the Eukaryotic 18S rRNA Endogenous Control (FAM/MGB probe, non-primer limited; Applied Biosystems #4333760F) in a 20 μL reaction, 10 μL TaqMan Fast Universal PCR Master Mix (Applied Biosystems #4364103) were mixed with 1 μL of primer-probe mix and 100 ng of cDNA template and subjected to the following cycling conditions: 50°C for 2 min; 95°C for 20 sec; followed by 40 cycles at 95°C for 3 s., and 55°C for 30 s.

### Analysis of library deletion enrichment

DVG RNA was extracted using the Quick-RNA Viral Kit (Zymo Research, USA). Amplicon libraries were prepared with the ARTIC V3 multiplex PCR protocol for SARS-CoV-2 (Integrate DNA Technologies, USA) and sequenced as paired-end reads on an Illumina MiniSeq platform. Raw reads were quality-filtered with fastp v0.23.2 [34], mapped to the DVG reference (DI) sequence using BBMap v.38.96 [35], and then deduplicated with samtools’ rmdup command [36]. The BAM file was converted to a readable tabular alignment format using samtools’ view command. To measure the encapsidation dependency of each position of the 667-nt PPS, we counted the number of DVG variants present in samples after 24 h. To derive a smooth function of the encapsidation dependency across the entire 667-nt PPS sequence, we took the average of the abundance of junctions within a span of 40 nucleotides on each side or each position of the PPS.

### Animal experimentation

Exposure of golden hamsters (*Mesocricetus auratus* (Waterhouse, 1839)) to SARS-CoV-2 was performed at the Integrated Research Facility at Fort Detrick (IRF-Frederick), National Institute for Allergy and Infectious Diseases (NIAID), Division of Clinical Research (DCR), National Institutes of Health (NIH). The IRF-Frederick is accredited by the Association for Assessment and Accreditation of Laboratory Animal Care (AAALAC; 000777), approved for Laboratory Animal Welfare by the Public Health Service (PHS; D16-00602), and registered with the United States Department of Agriculture (USDA; 51-F-0016). Experimental procedures were approved by the NIAID DCR Animal Care and Use Committee (animal study protocol #IRF-070E) and followed the recommendations provided in The Guide for the Care and Use of Laboratory Animals [37], the American Veterinary Medical Association (AVMA) guidelines for the euthanasia of animals [38], and the ARRIVE (Animal Research: Reporting *In Vivo* Experiments) guidelines 2.0 [39]. Work with infectious SARS-CoV-2 at the NIH NIAID DCR IRF-Frederick was approved by the Institutional Biosafety Committee (IBC).

Six 5-week-old golden hamsters were assigned per group. Hamsters in each infected group were exposed to 10^6^ PFU of SARS-CoV-2 per animal through intranasal delivery at Day 0. Hamsters in the uninfected group were exposed to mock control consisting of PBS. 14 h post-exposure, hamsters were given an intranasal dose of 60 μl of hPBAE-RNA polyplexes in PBS containing 1 mg of RNA. At day 5, hamsters were anesthetized via 4% isoflurane in an induction box and euthanised. Total RNA was extracted from hamster lungs using the KingFisher Flex automated extraction system (Thermo Fisher Scientific) in combination with the MagMAX RNA Isolation Kit, following the manufacturer’s recommended procedures. Tissues were first homogenized in lysis buffer to disrupt cells and release nucleic acids. Homogenates were then processed on the KingFisher Flex, which uses magnetic bead–based capture to bind RNA, followed by sequential wash steps to remove contaminants. Purified RNA was eluted in RNase-free water and stored at −80°C until downstream analysis.

## ACKNOWLEDGEMENTS

We acknowledge the Huck Institutes’ Genomics Core Facility (RRID:SCR_023645) for sequencing and we thank the National Institutes of Health (NIH) National Institute of Allergy and Infectious Diseases (NIAID) Division of Clinical Research (DCR) Integrated Research Facility at Fort Detrick (IRF-Frederick) Comparative Medicine and Clinical Core staff for successful implementation of the animal studies.

## AUTHOR CONTRIBUTIONS

SY: synthetised and validated constructs, analyzed data

AG: performed in vitro experiments, analyzed data, revised the manuscript

JA: performed in vivo experiments and analyzed data

MB: performed bioinformatic analysis

PJ: analyzed in vitro data

GW: designed and supervised in vivo experiments

JHK: supervised in vivo experiments, revised the manuscript

SVK: designed and supervised in vitro experiments, revised the manuscript

MA: supervised research, designed constructs and experiments, analyzed and interpreted data, performed comparative and RNA structure analysis, and wrote the manuscript

## SUPPLEMENTARY FIGURES

**Supplementary Fig. 1.**
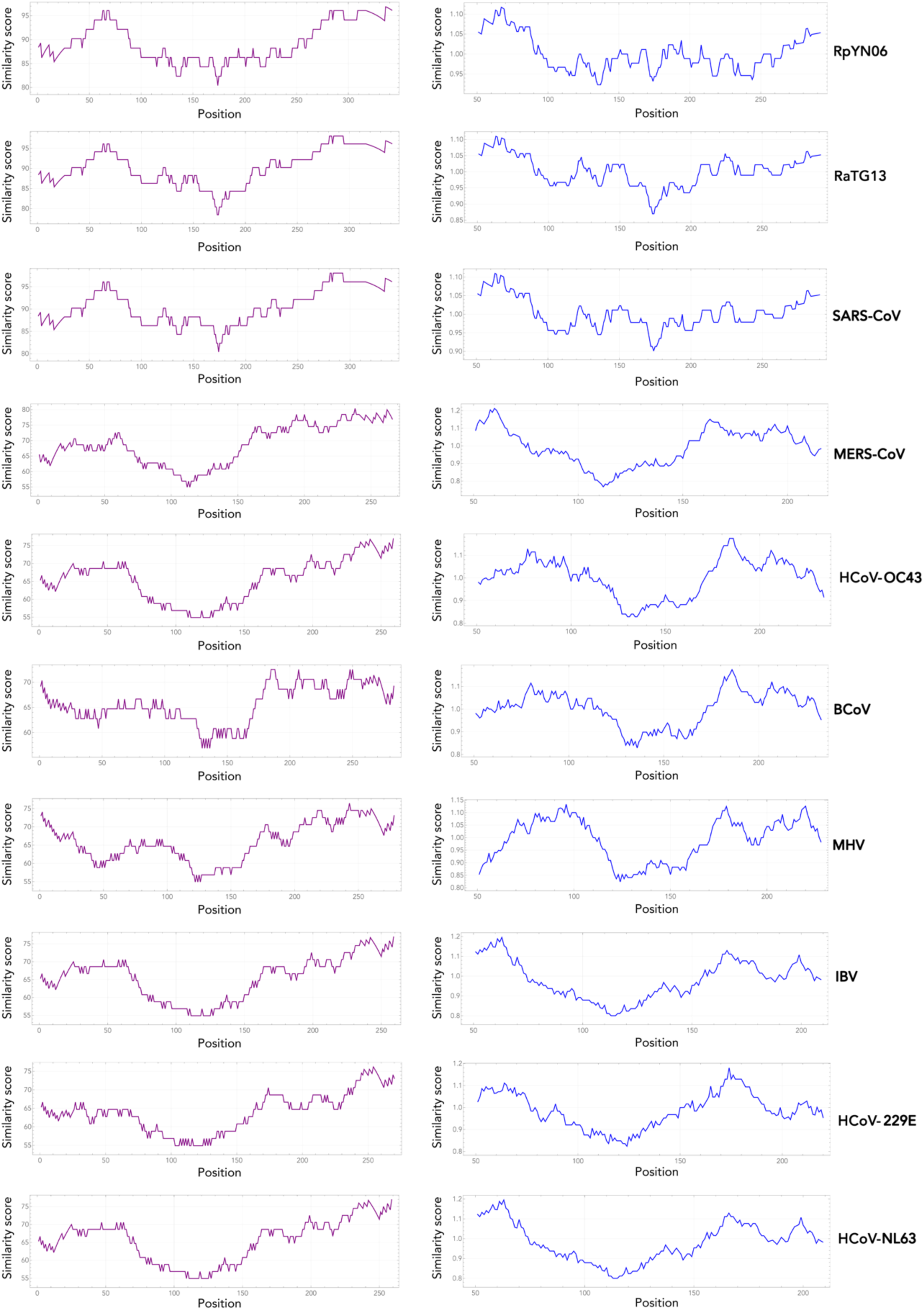
Sequence similarity of SARS-CoV-2 nonstructural protein 15 (Nsp15) to coronavirus homologs. Purple (left): absolute similarity. Blue (right): local similarity, relative to flanking regions.

**Supplementary Fig. 2.**
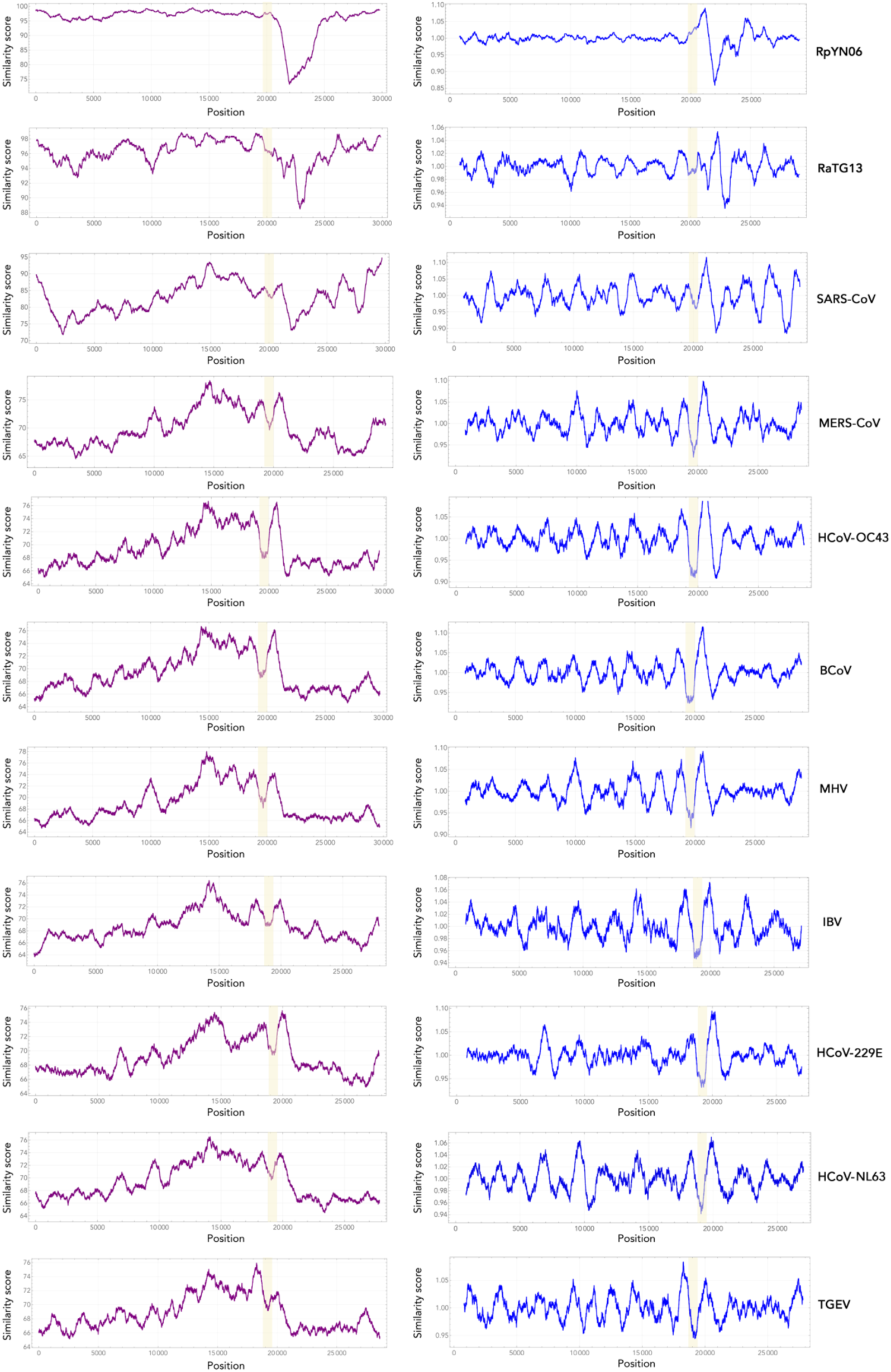
Sequence similarity of SARS-CoV-2 genome to the genomes of other coronaviruses. Purple (left): absolute similarity. Blue (right): local similarity, relative to flanking regions. The 667-nt PPS is highlighted in yellow.

**Supplementary Fig. 3.**
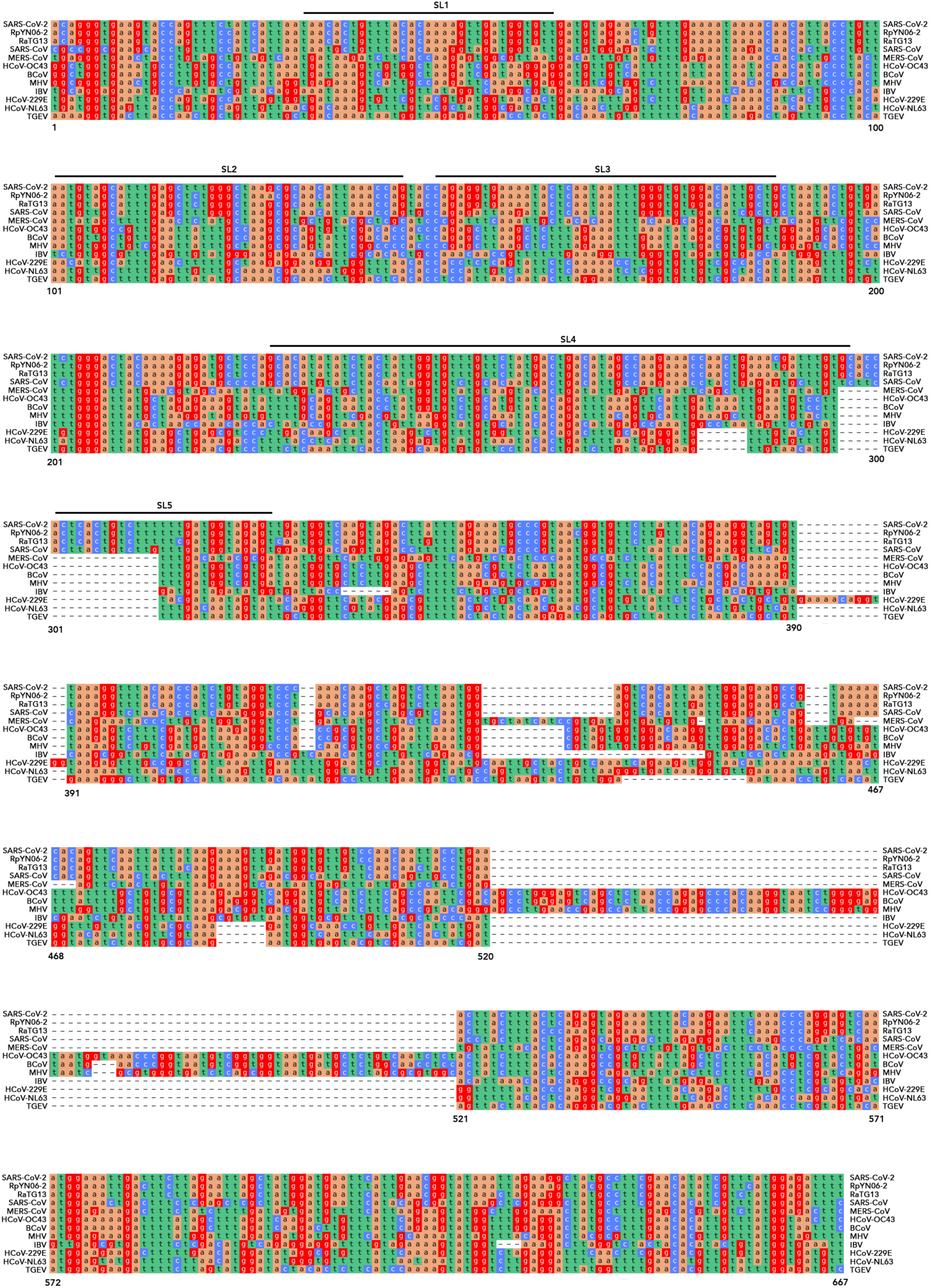
Alignment of the 667-nt PPS of SARS-CoV-2 with the genome of other coronaviruses. Regions SL1-SL5 refer to stem loops in Supplementary Fig. 7−8.

**Supplementary Fig. 4.**
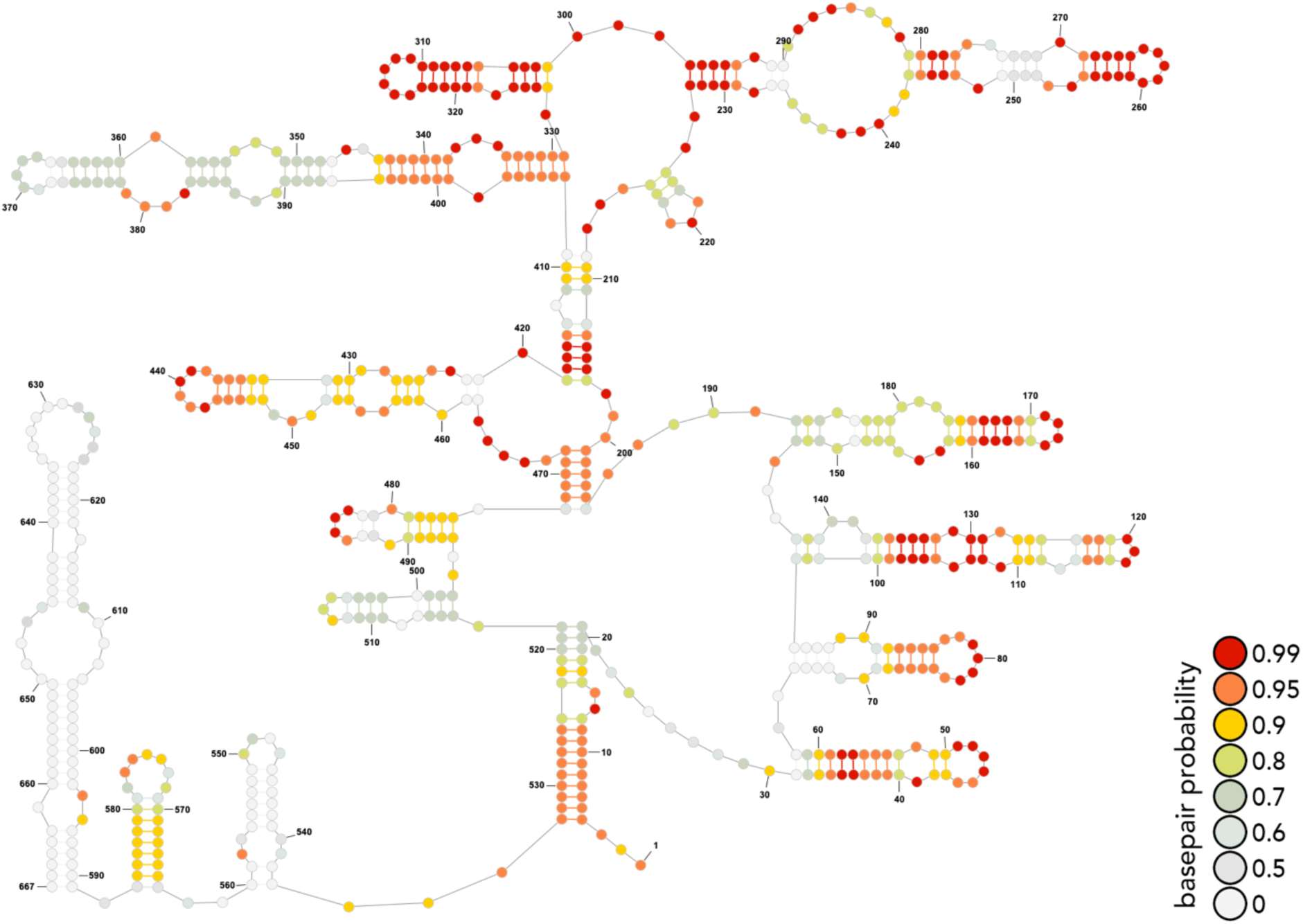
Predicted secondary structure of the 667-nt PPS.

**Supplementary Fig. 5.**
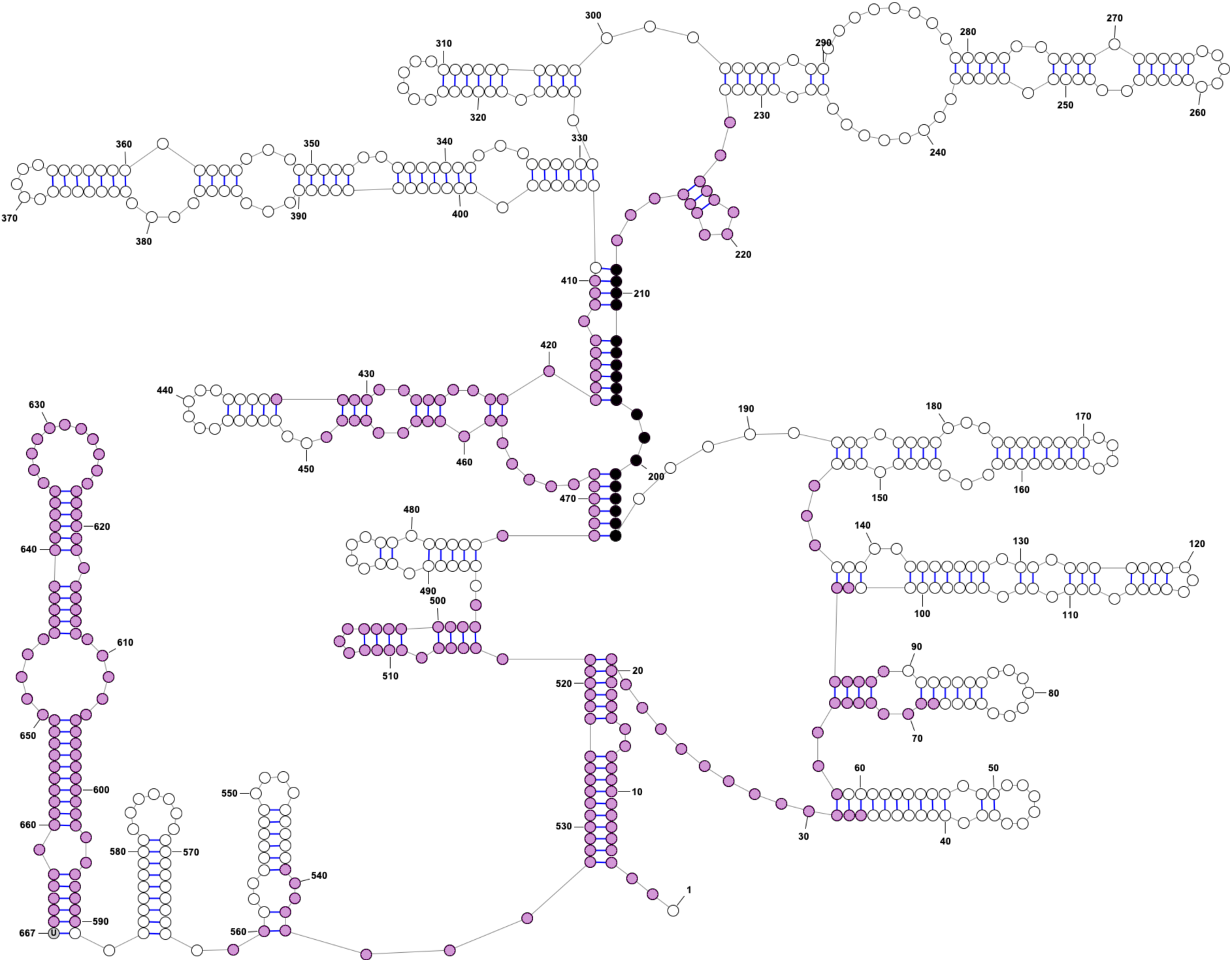
Secondary structures within the 667-nt PPS disrupted by deleting nucleotides 194-212. Shown is the predicted secondary structure of the 667-nt PPS. Deleting nucleotides 194−212 (black) disrupts the predicted secondary structures depicted in purple. See also Supplementary Fig. 6.

**Supplementary Fig. 6.**
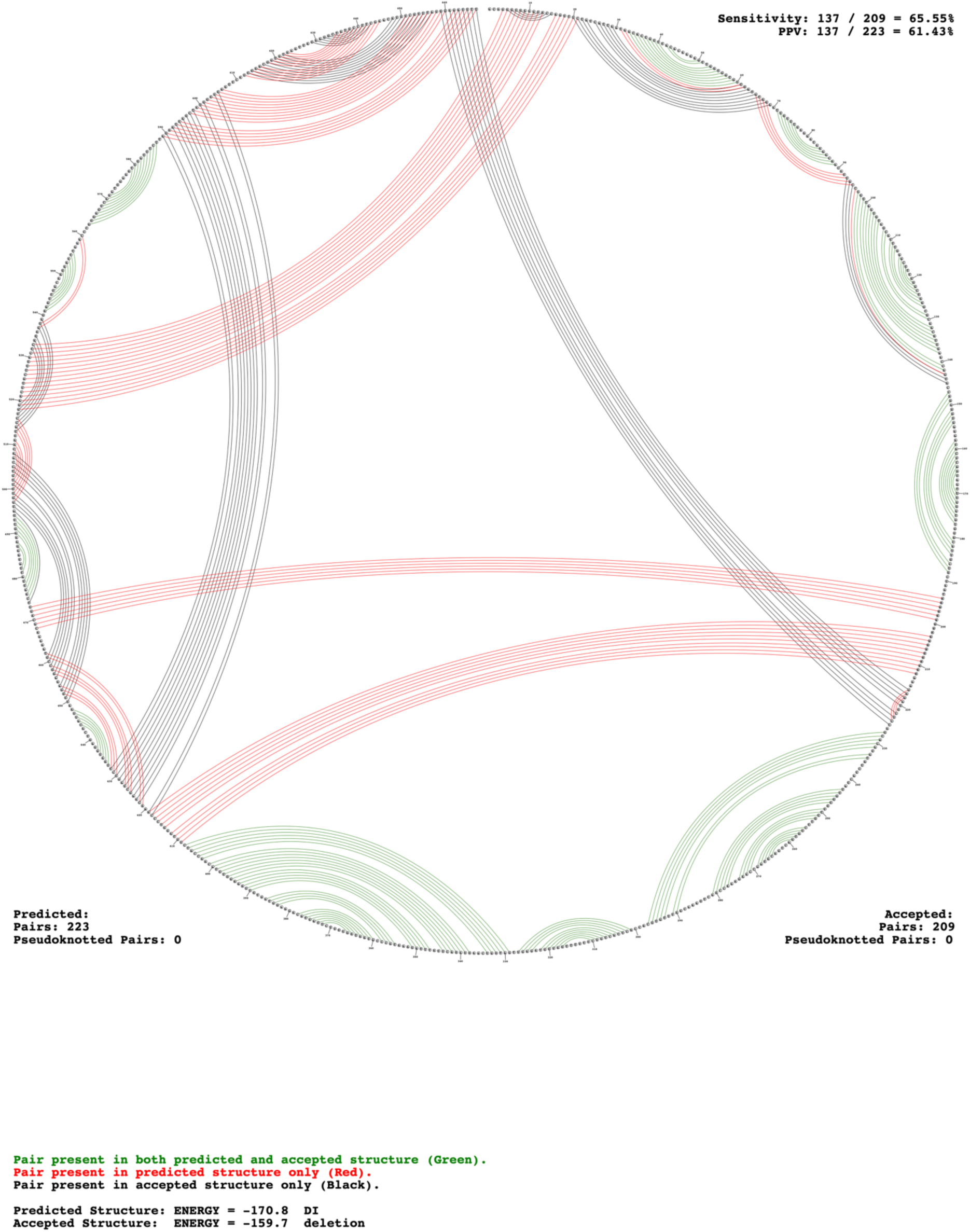
Secondary structures within the 667-nt PPS disrupted by deleting nucleotides 194-212. Same as Supplementary Fig. 5 but drawn as a circle plot. Pairings in green are predicted to occur in both sequences. Pairings in red are predicted to occur only in the 667-nt PPS. Pairings in black are predicted to occur only when nucleotides 194−212 are deleted.

**Supplementary Fig. 7.**
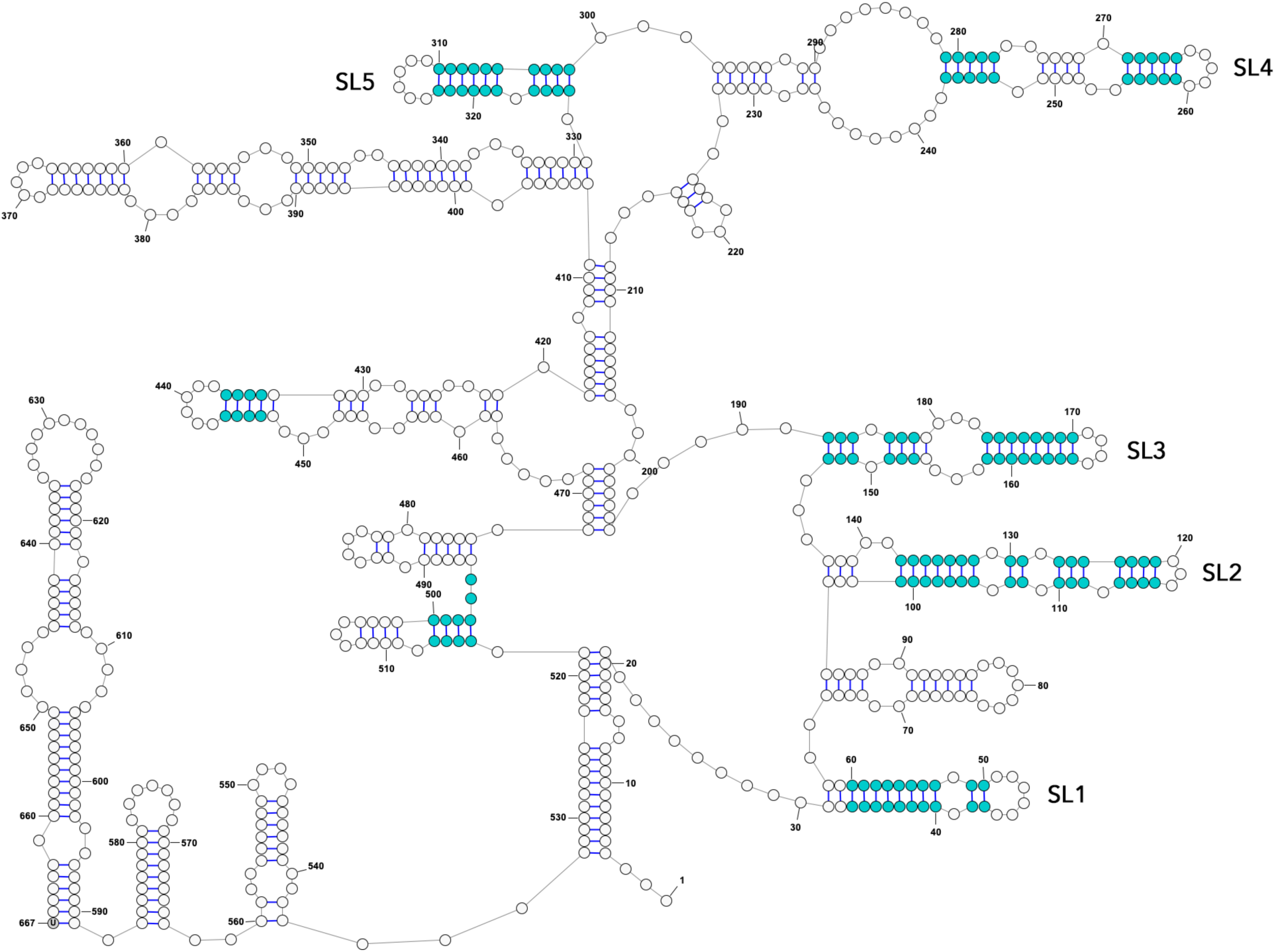
Predicted secondary structures within the 667-nt PPS that are also predicted in the homologous region of SARS-CoV. Shown is the predicted secondary structure of the 667-nt PPS. Base-pairs depicted in cyan are predicted to be shared between SARS-CoV-2 and SARS-CoV. See also Supplementary Fig. 8.

**Supplementary Fig. 8.**
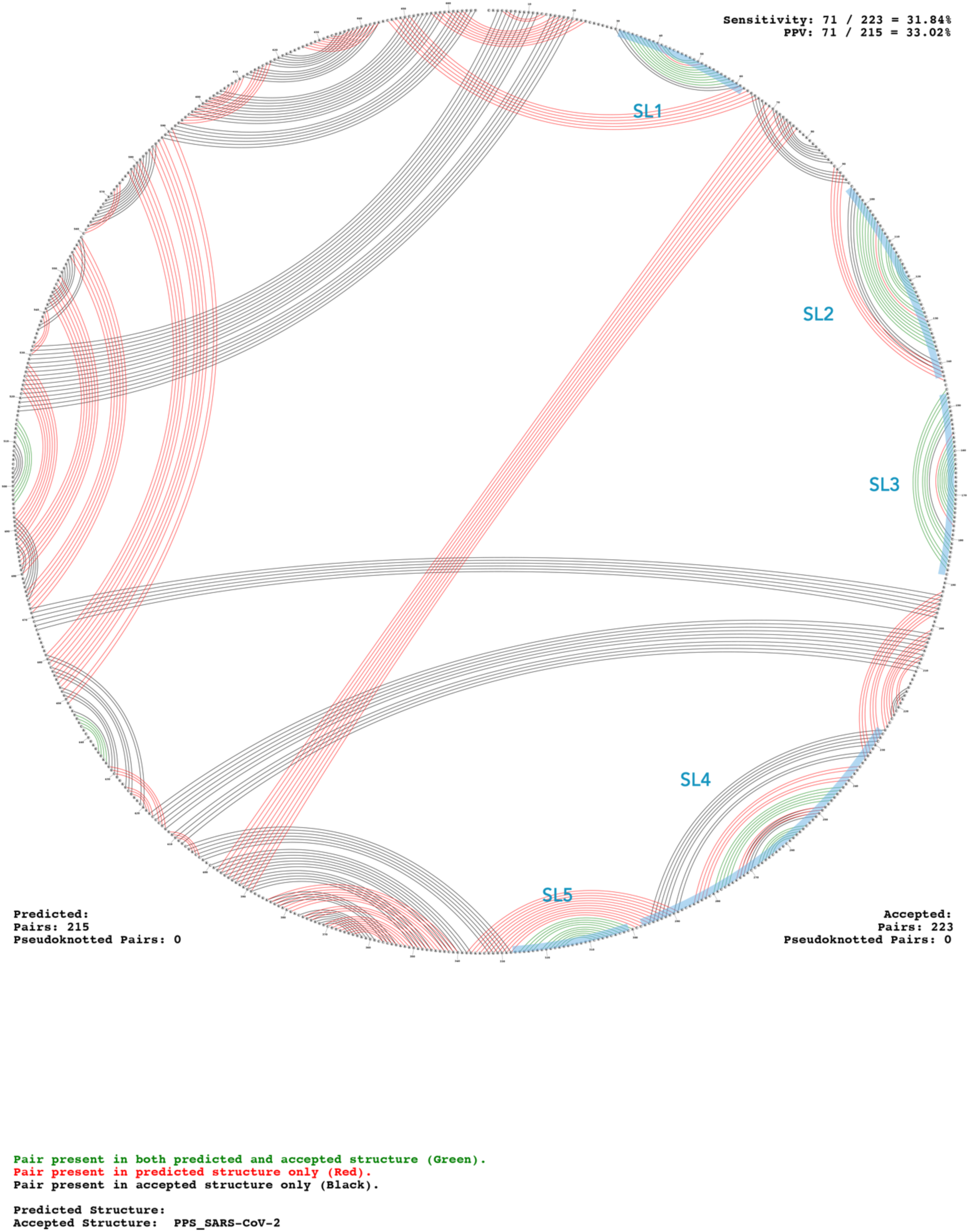
Predicted secondary structures within the 667-nt PPS that are also predicted in the homologous region of SARS-CoV. Same as Supplementary Fig. 7 but drawn as a circle plot. Predicted stem loops SL1−SL5 are highlighted in blue. Pairings in green are predicted to occur in both sequences. Pairings in red are predicted to occur only in SARS-CoV-2. Pairings in black are predicted to occur only in SARS-CoV.

**Supplementary Fig. 9.**
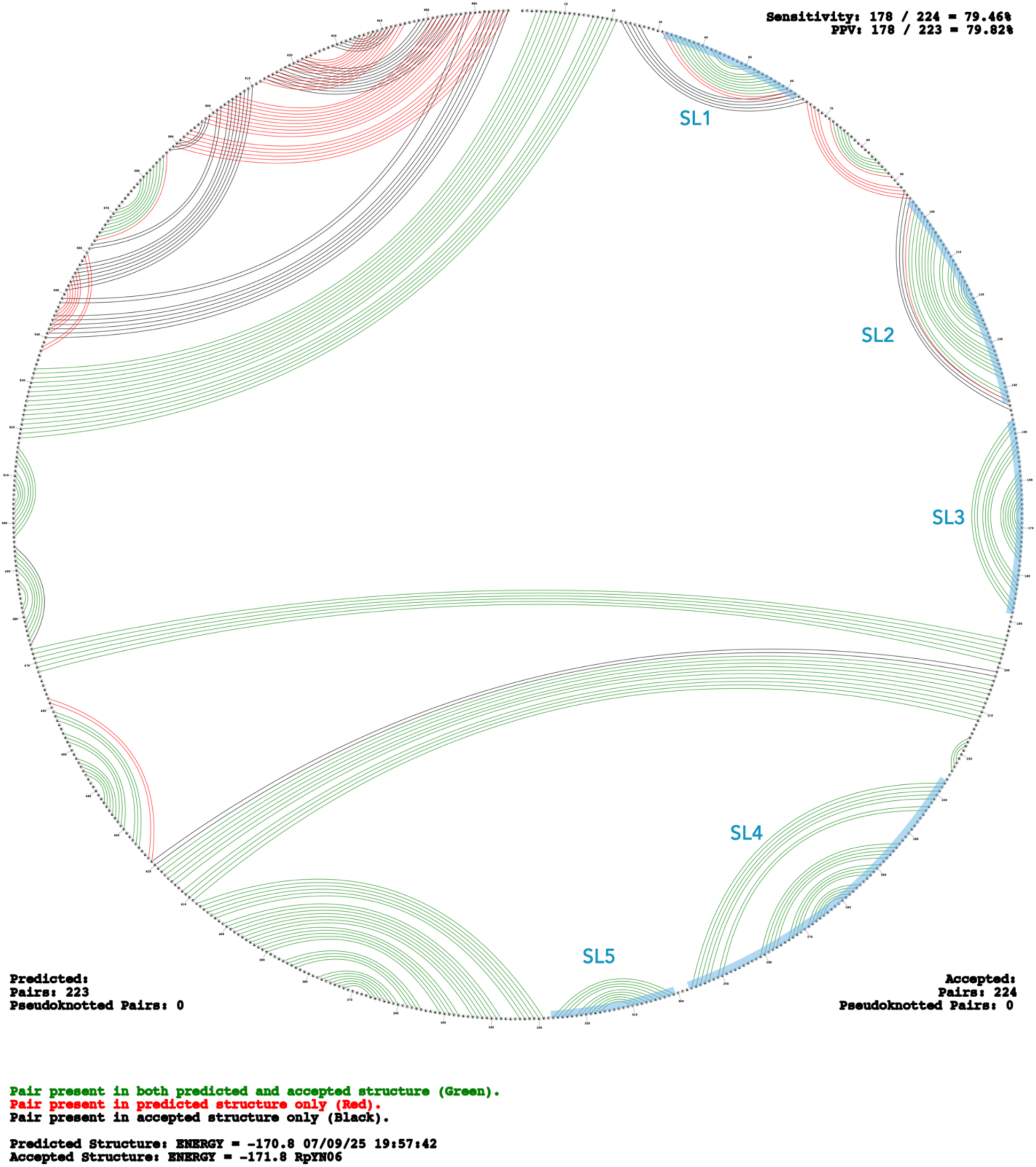
Predicted secondary structures within the 667-nt PPS that are also predicted in the homologous region of RpYN06. Predicted stem loops SL1−SL5 are highlighted in blue. Pairings in green are predicted to occur in both sequences. Pairings in red are predicted to occur only in SARS-CoV-2. Pairings in black are predicted to occur only in RpYN06.

**Supplementary Fig. 10.**
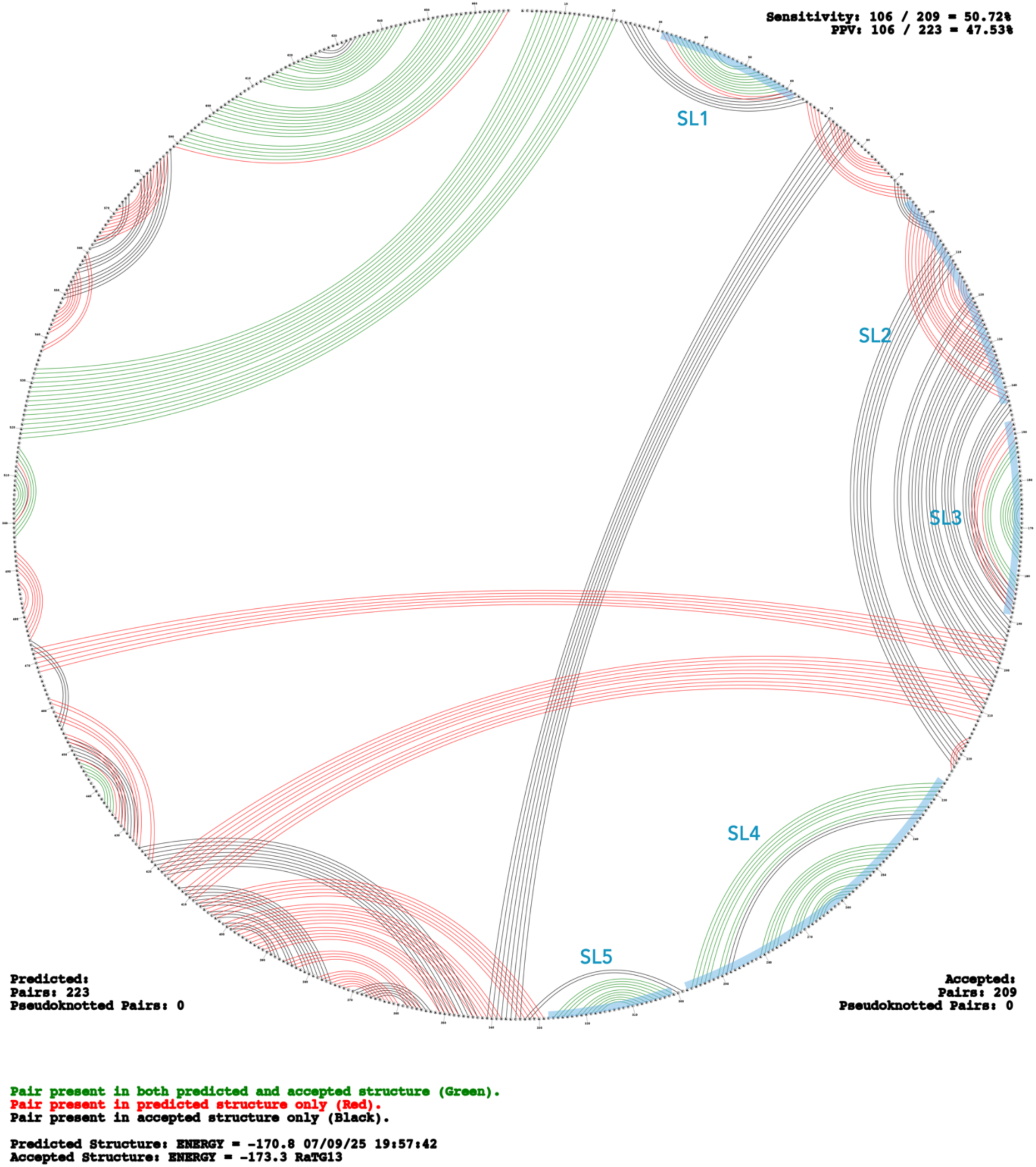
Predicted secondary structures within the 667-nt PPS that are also predicted in the homologous region of RaTG13. Predicted stem loops SL1−SL5 are highlighted in blue. Pairings in green are predicted to occur in both sequences. Pairings in red are predicted to occur only in SARS-CoV-2. Pairings in black are predicted to occur only in sarbecovirus RaTG13.

**Supplementary Fig. 11.**
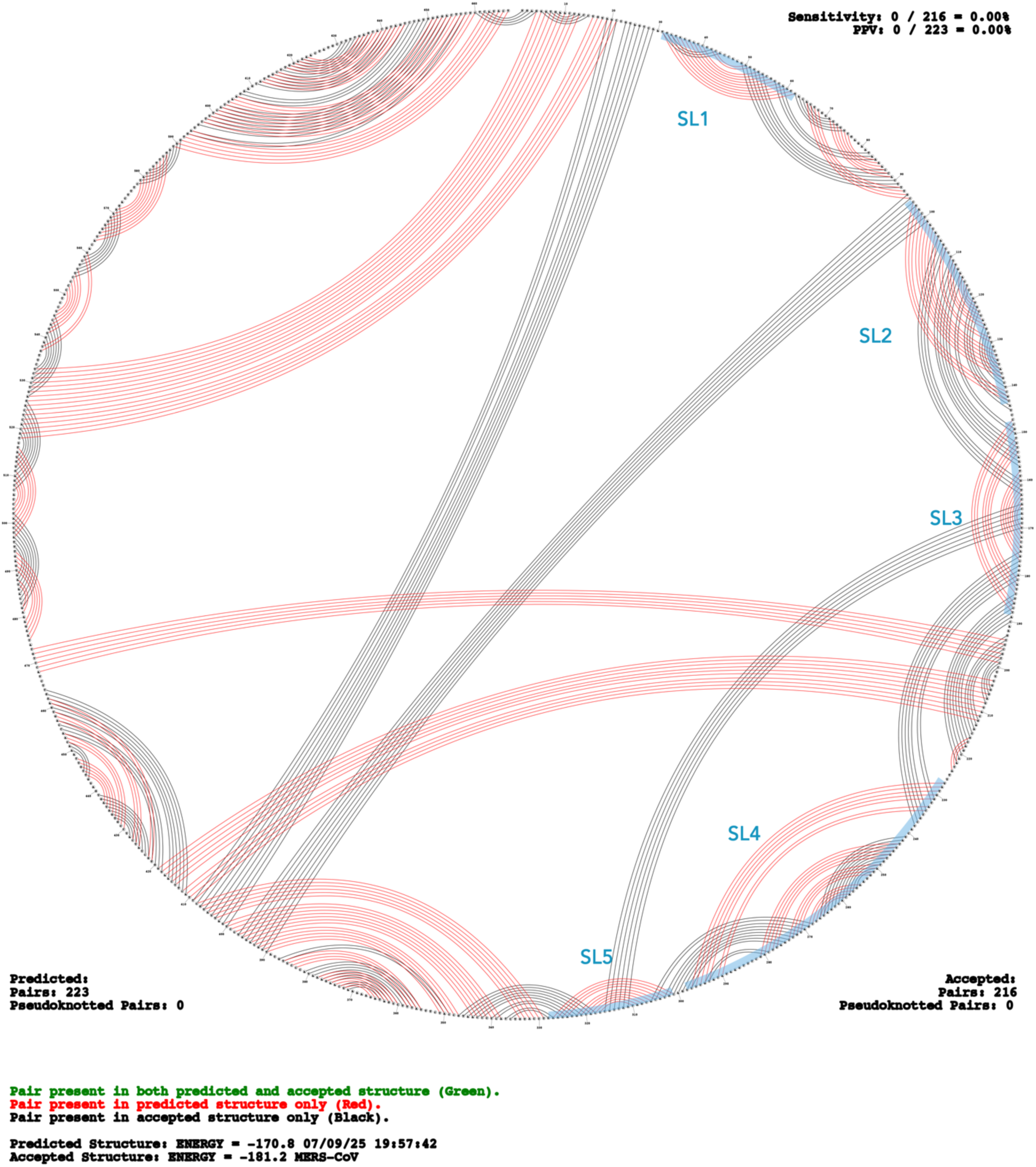
Predicted secondary structures within the 667-nt PPS that are also predicted in the homologous region of MERS-CoV. Predicted stem loops SL1−SL5 are highlighted in blue. Pairings in green are predicted to occur in both sequences. Pairings in red are predicted to occur only in SARS-CoV-2. Pairings in black are predicted to occur only in MERS-CoV.

**Supplementary Fig. 12.**
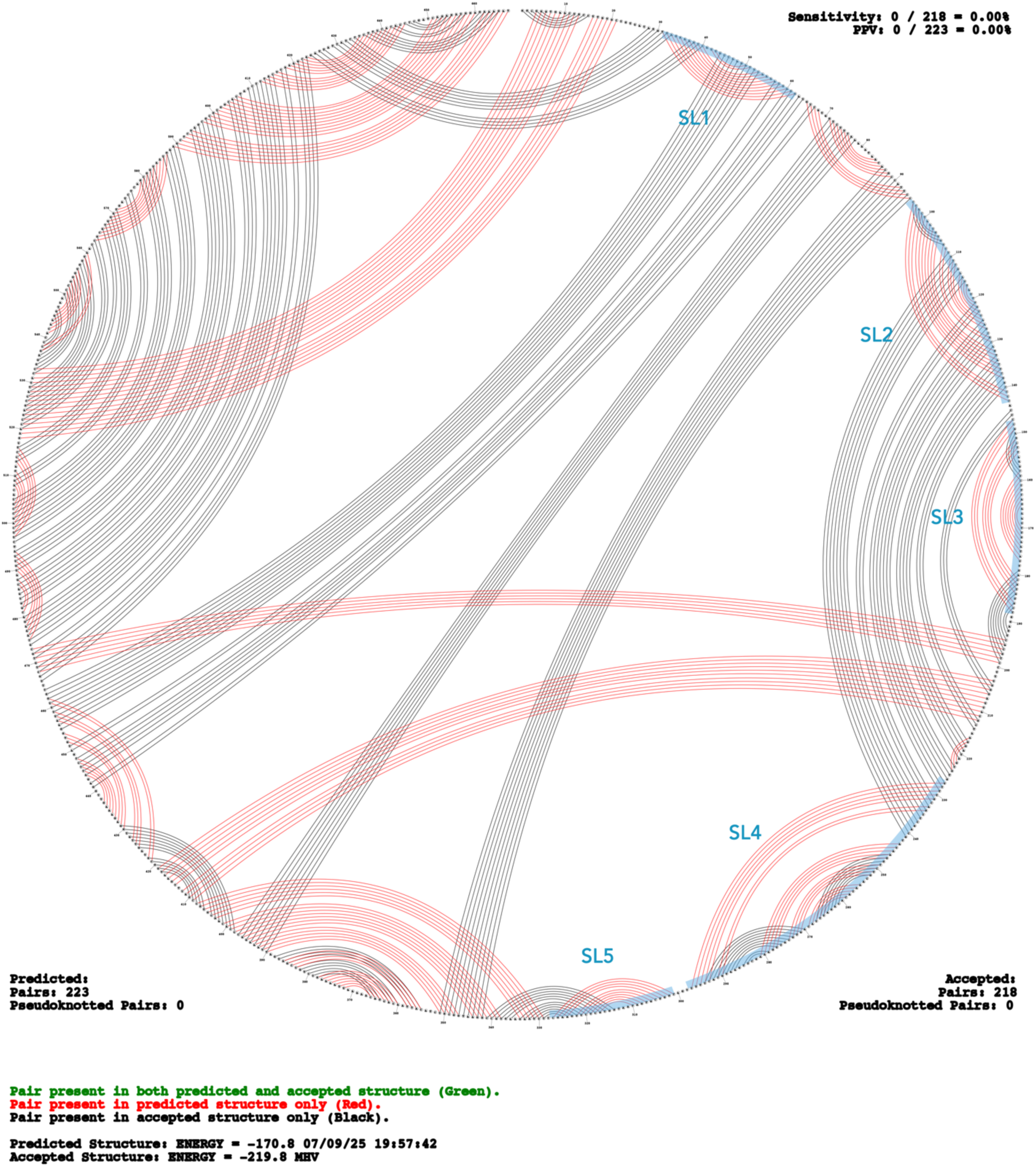
Predicted secondary structures within the 667-nt PPS that are also predicted in the homologous region of MHV. Predicted stem loops SL1−SL5 are highlighted in blue. Pairings in green are predicted to occur in both sequences. Pairings in red are predicted to occur only in SARS-CoV-2. Pairings in black are predicted to occur only in MHV.

**Supplementary Fig. 13.**
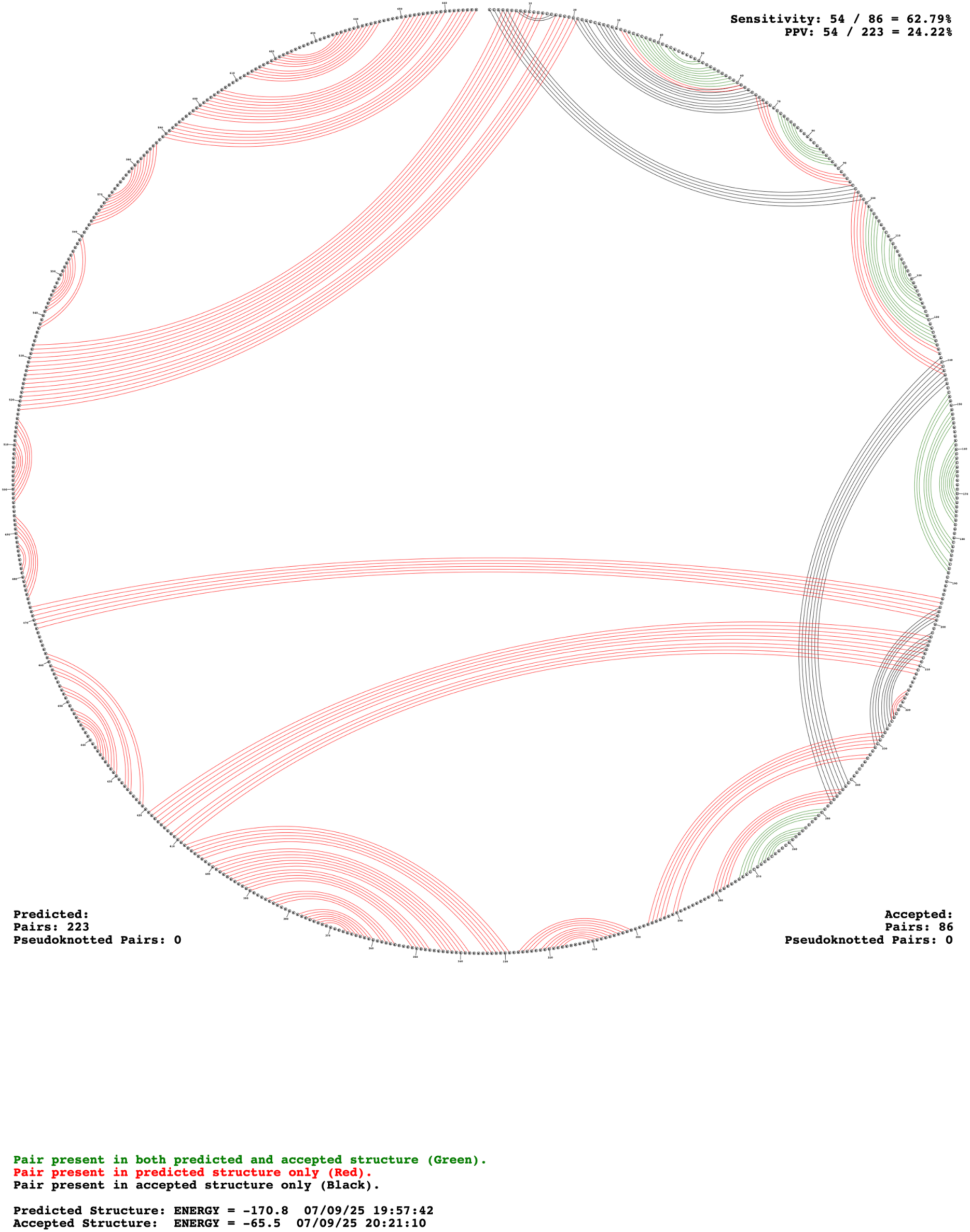
Predicted secondary structures within the 667-nt PPS that are also predicted to occur in DVG F6. Pairings in green are predicted to occur in both sequences. Pairings in red are predicted to occur only in the 667-nt PPS. Pairings in black are predicted to occur only in DVG F6.

**Supplementary Fig. 14.**
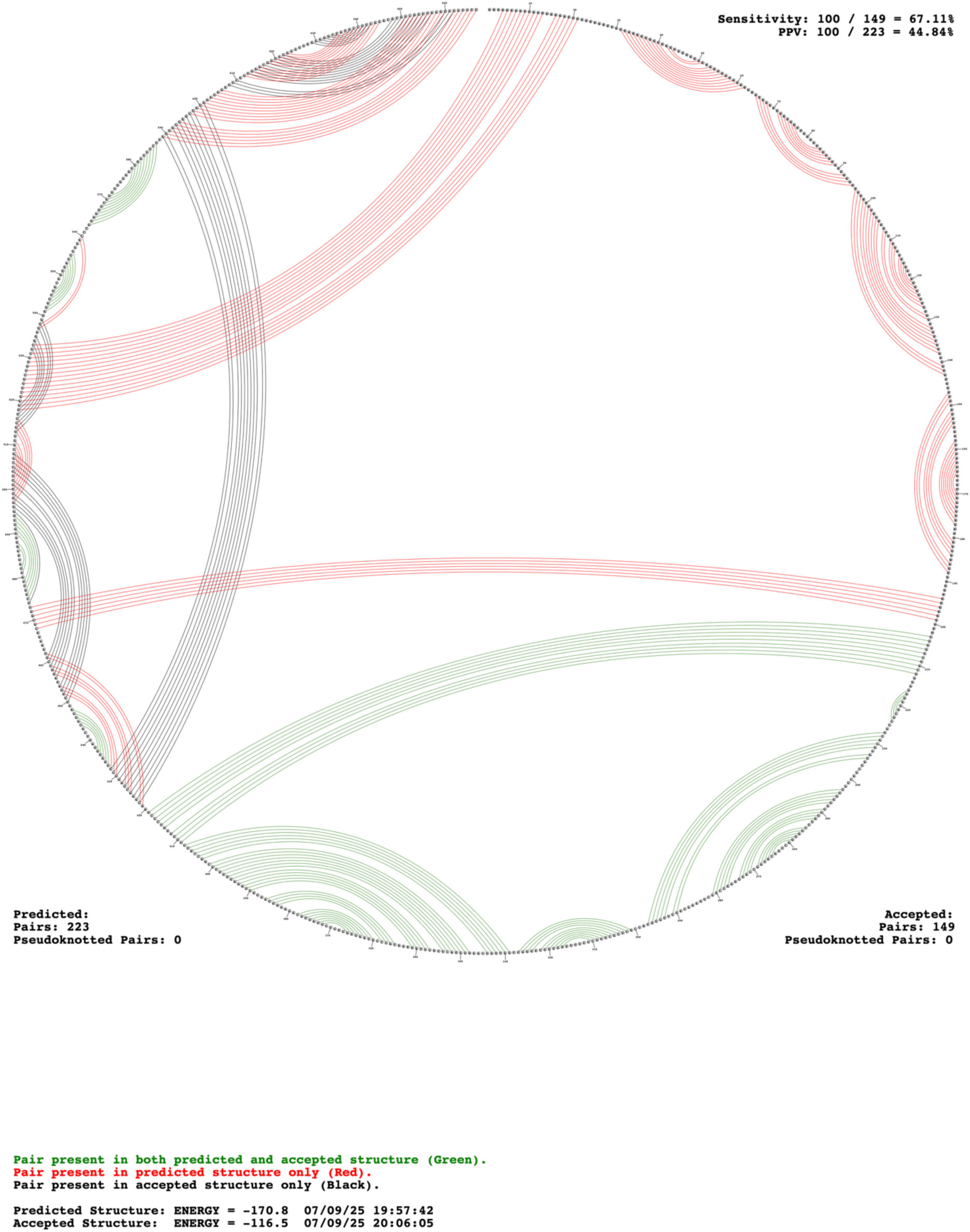
Predicted secondary structures within the 667-nt PPS that are also predicted to occur in DVG F7. Pairings in green are predicted to occur in both sequences. Pairings in red are predicted to occur only in the 667-nt PPS. Pairings in black are predicted to occur only in DVG F7.

**Supplementary Fig. 15.**
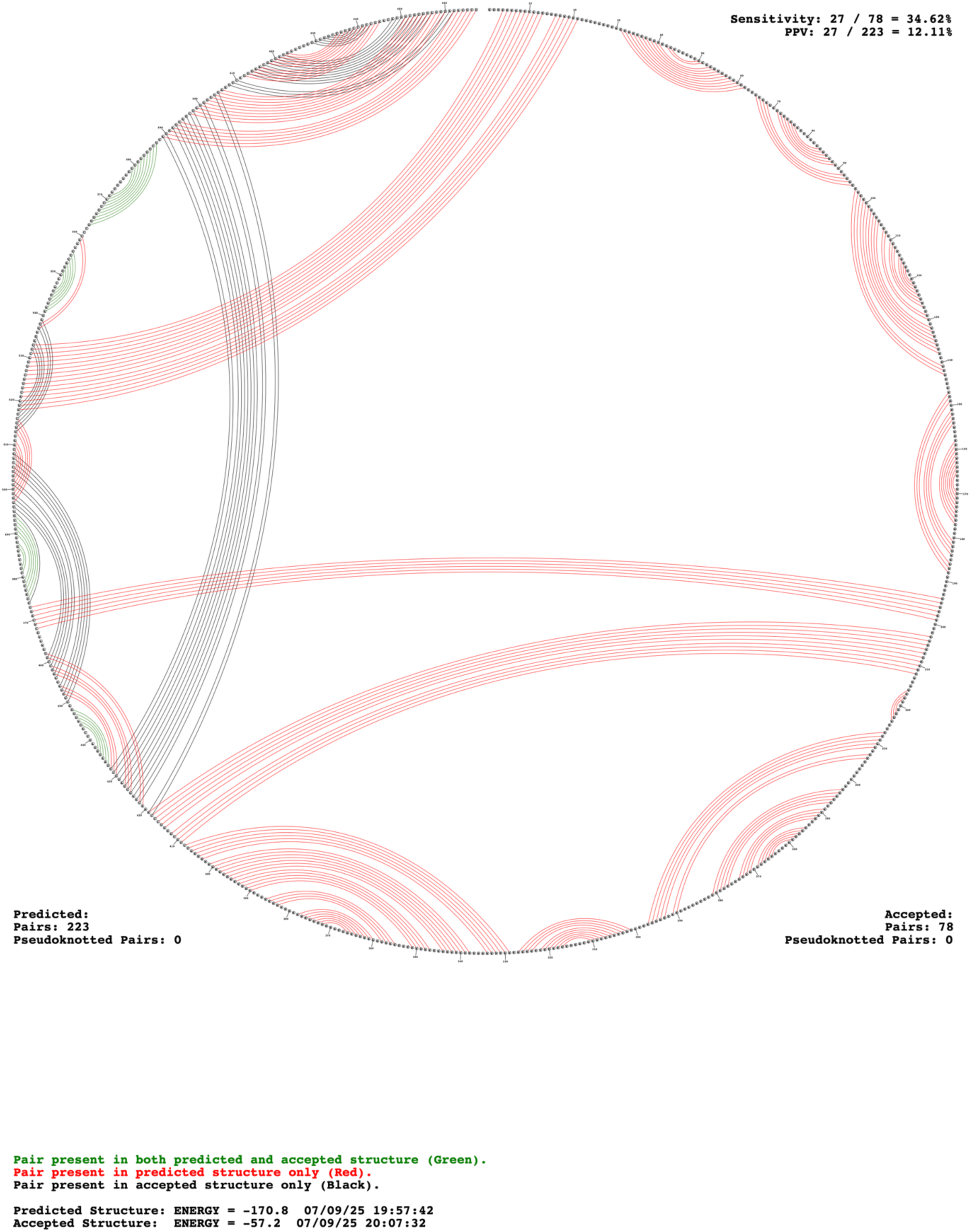
Predicted secondary structures within the 667-nt PPS that are also predicted to occur in sequence PS9. Pairings in green are predicted to occur in both sequences. Pairings in red are predicted to occur only in the 667-nt PPS. Pairings in black are predicted to occur only in PS9.

**Supplementary Fig. 16.**
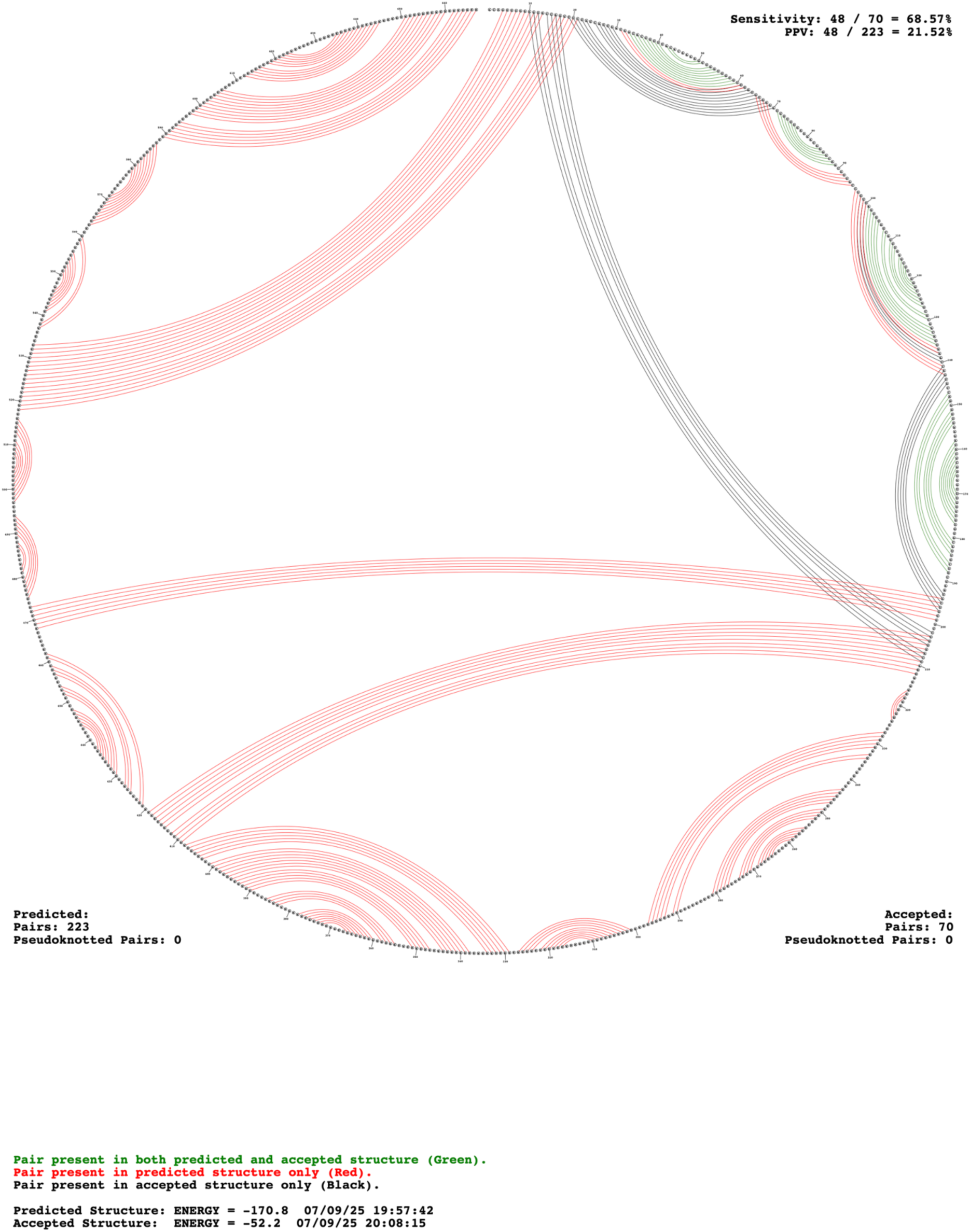
Predicted secondary structures within the 667-nt PPS that are also predicted to occur in sequence PS576. Pairings in green are predicted to occur in both sequences. Pairings in red are predicted to occur only in the 667-nt PPS. Pairings in black are predicted to occur only in PS576.

**Supplementary Fig. 17.**
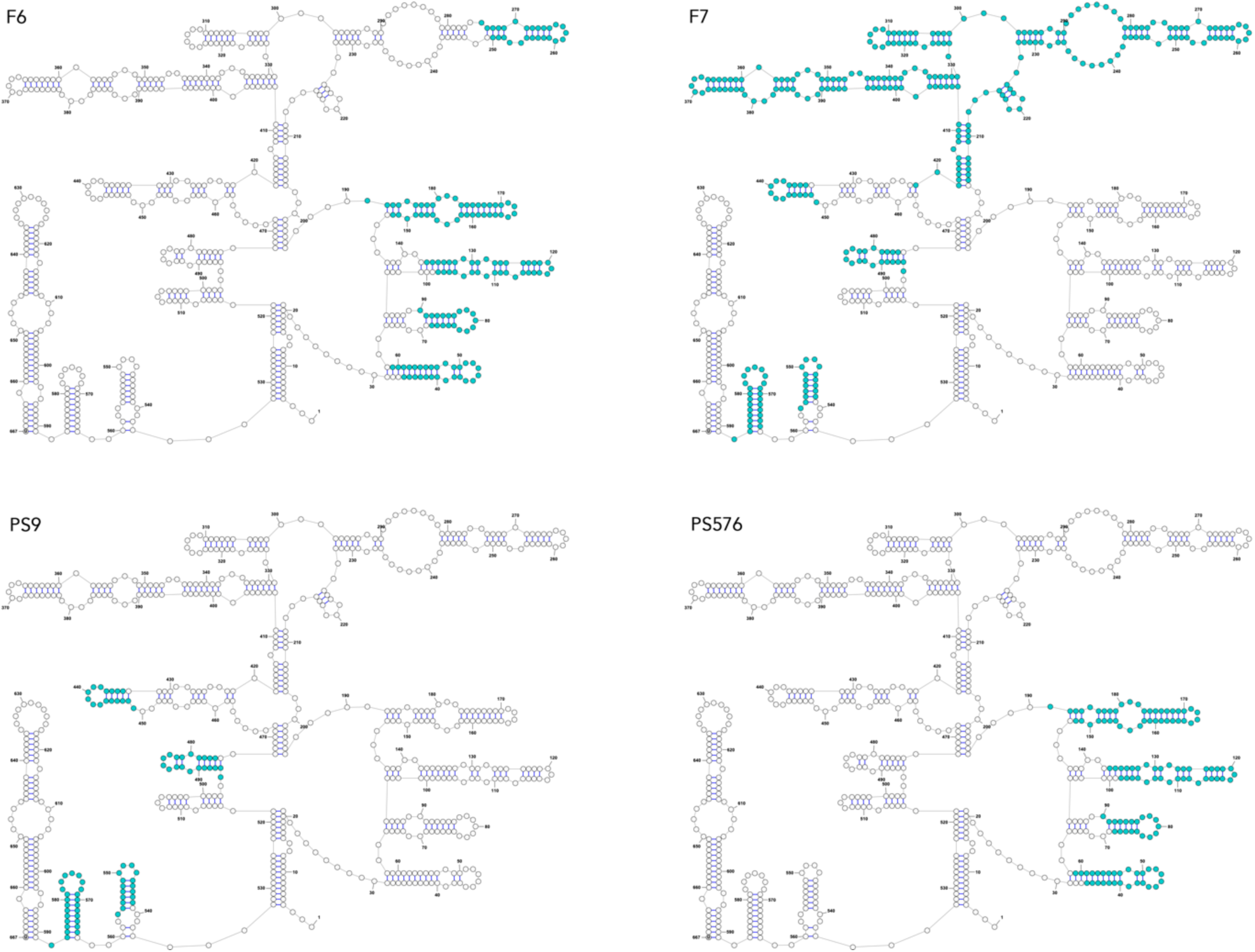
Predicted secondary structures within the 667-nt PPS that are also predicted to occur in sequences F6, F7, PS9 and PS576. Shown is the predicted secondary structure of the 667-nt PPS. The parts in cyan are secondary structures also predicted to occur in F6, F7, PS9 and PS576 (see also the corresponding circle plots in Supplementary Fig. 9−12).

**Supplementary Fig. 18.**
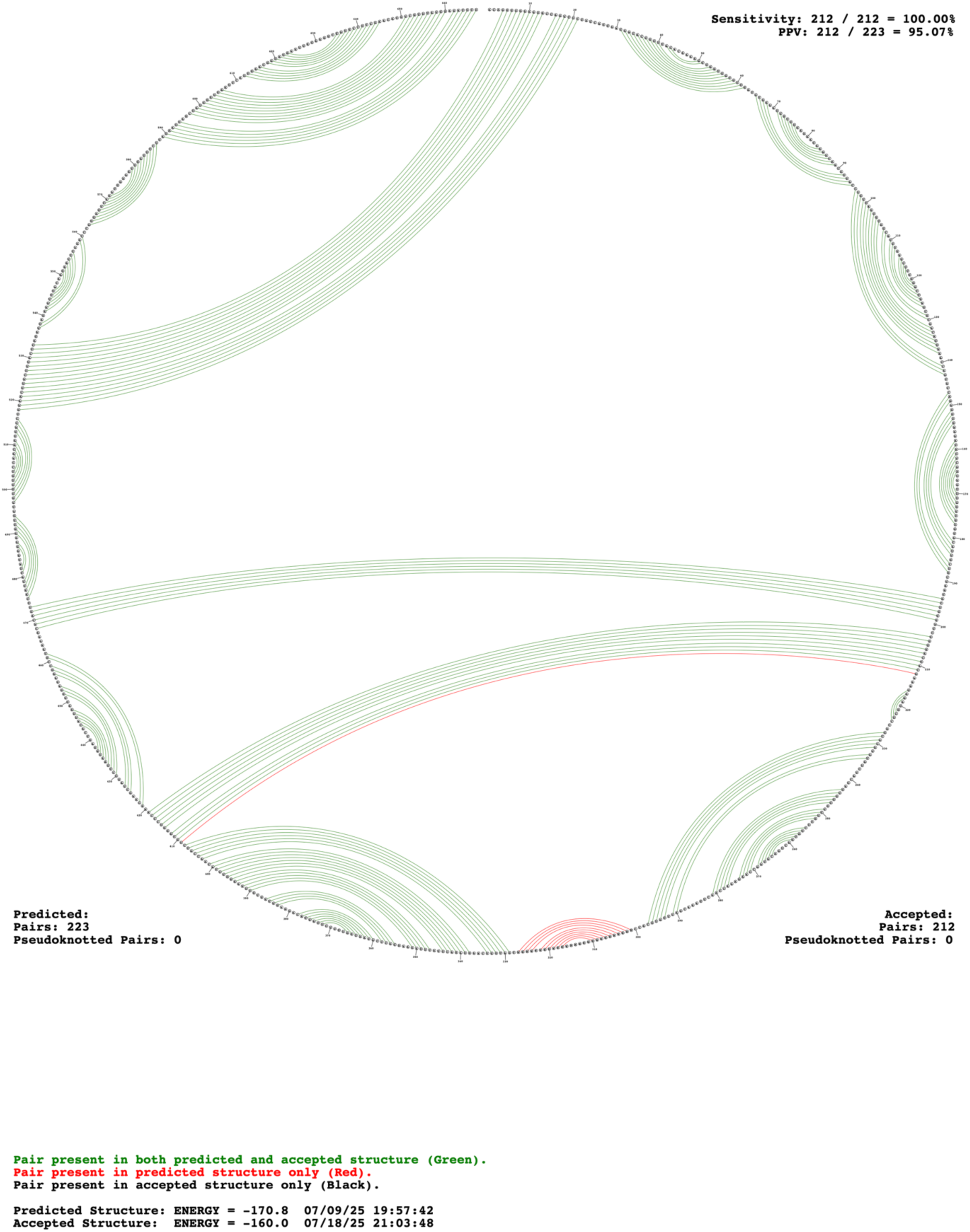
Predicted secondary structures within the 667-nt PPS that are predicted to be disrupted in DIΔ1. Pairings in green are predicted to occur in both sequences. Pairings in red are predicted to occur only in the 667-nt PPS. Pairings in black are predicted to occur only in DIΔ1.

**Supplementary Fig. 19.**
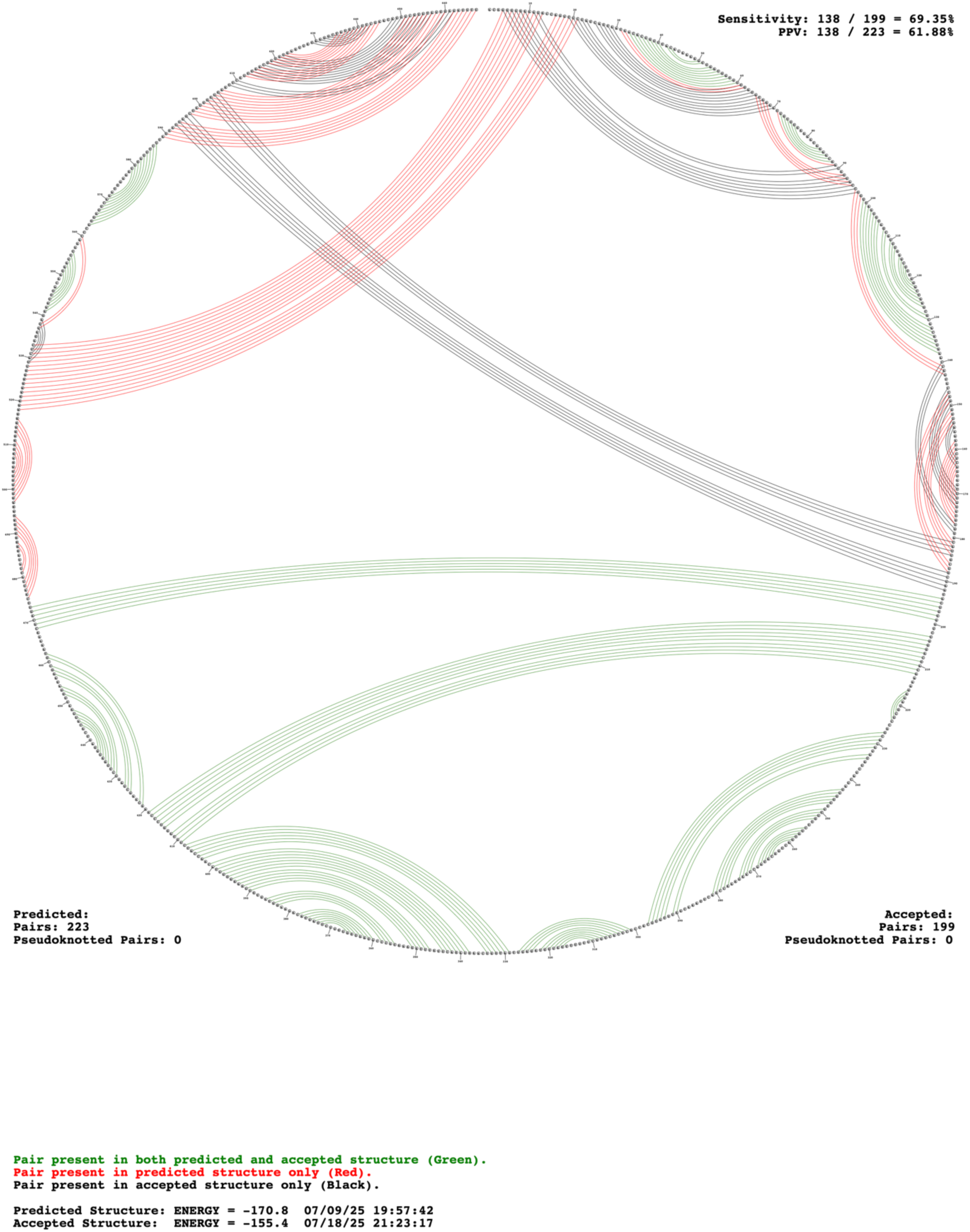
Predicted secondary structures within the 667-nt PPS that are predicted to be disrupted in DIΔ2. Pairings in green are predicted to occur in both sequences. Pairings in red are predicted to occur only in the 667-nt PPS. Pairings in black are predicted to occur only in DIΔ2.

**Supplementary Fig. 20.**
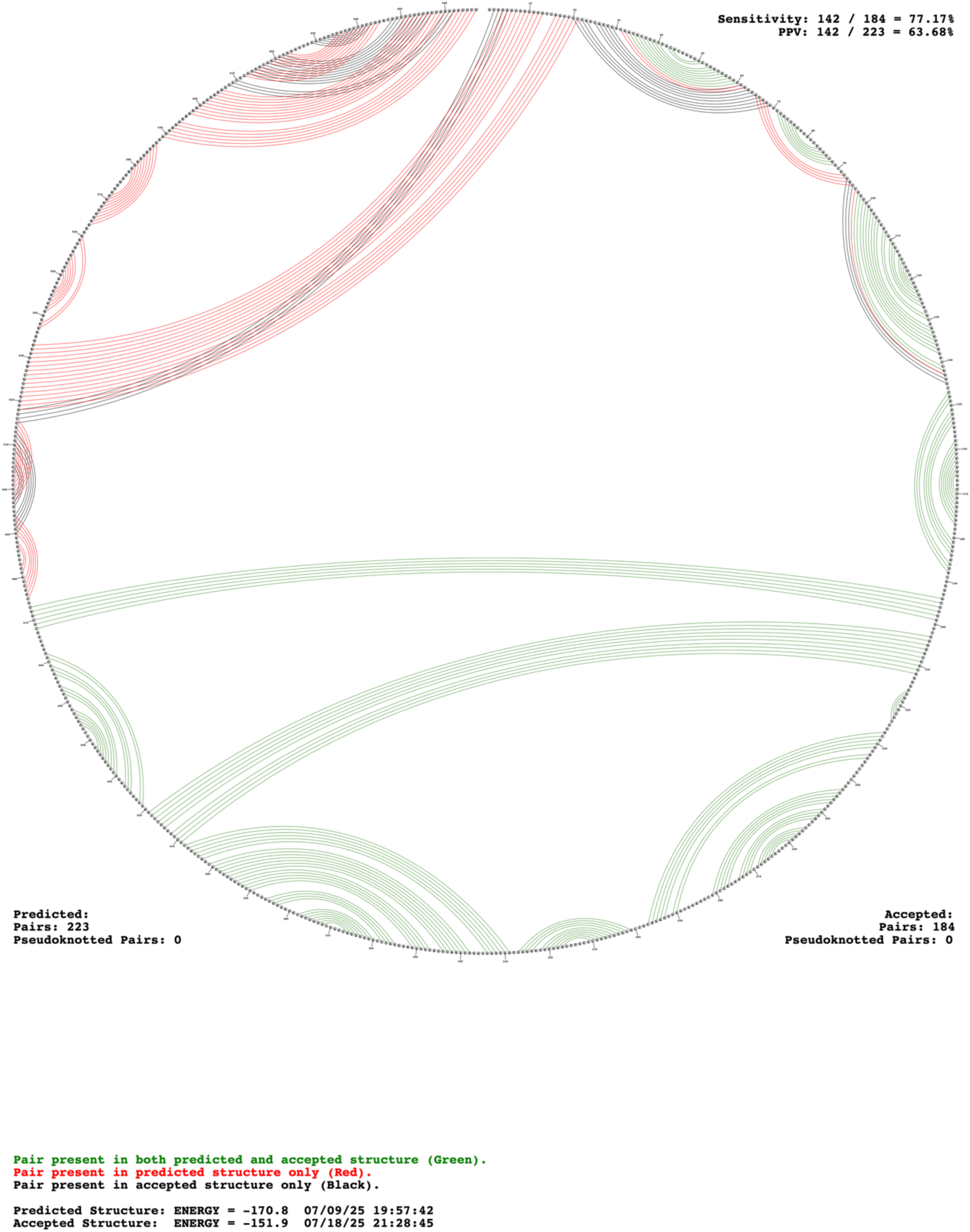
Predicted secondary structures within the 667-nt PPS that are predicted to be disrupted in DIΔ3. Pairings in green are predicted to occur in both sequences. Pairings in red are predicted to occur only in the 667-nt PPS. Pairings in black are predicted to occur only in DIΔ3.

**Supplementary Fig. 21.**
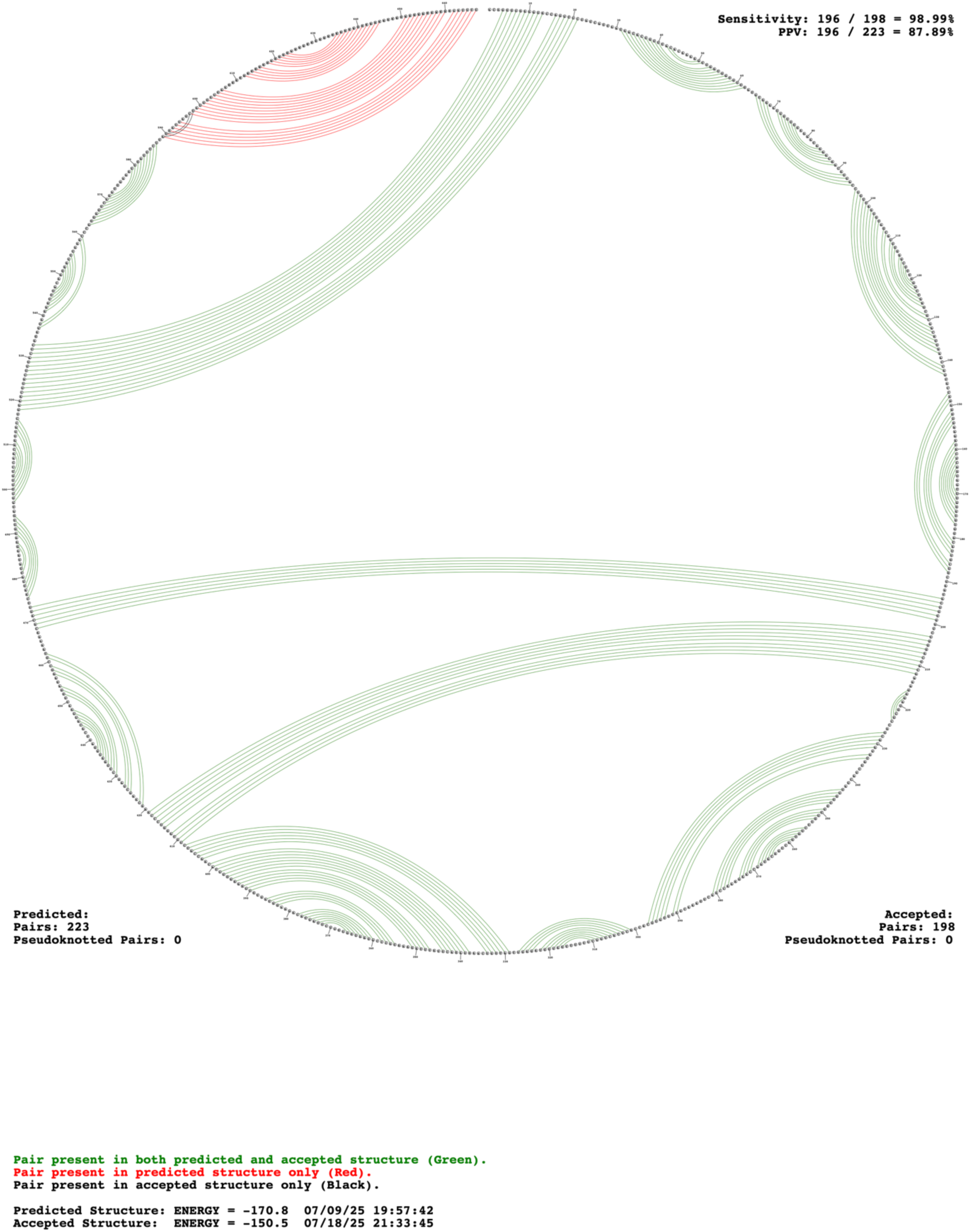
Predicted secondary structures within the 667-nt PPS that are predicted to be disrupted in DIΔ4. Pairings in green are predicted to occur in both sequences. Pairings in red are predicted to occur only in the 667-nt PPS. Pairings in black are predicted to occur only in DIΔ4.

